# Pathogenome and Plasmid-Borne Antimicrobial Resistance Phenotypes in a Multidrug-Resistant O111:H8 Shiga Toxin-Producing *Escherichia coli* Strain

**DOI:** 10.64898/2026.09.16.752176

**Authors:** Irvin Rivera, Anwar A. Kalalah, Sara S.K. Koenig, Mariana Sainz Garcia, Jacob A. Alford, Armando L. Rodriguez, Joseph M. Bosilevac, Mark Eppinger

**Author notes:** Corresponding author: Mark Eppinger, PhD, MS, Department of Molecular Microbiology and Immunology (MMI) & South Texas Center for Emerging Infectious Diseases (STCEID), The University of Texas at San Antonio (UTSA), One UTSA Circle, San Antonio, TX, 78249-0600.

## Abstract

Plasmids contribute to virulence and antimicrobial resistance in Shiga toxin-producing *Escherichia coli* (STEC). Here, the mobility, gene content, and evolutionary context of four plasmids carried by an O111:H8 STEC strain designated UTAK-22: pUTAK-22.1-MDR, pUTAK-22.2-MDR, pUTAK-22.3-P1, and pUTAK-22.4-pO111 were characterized. Conjugation experiments, phenotype-based selective readouts, and comparative genomics together showed that pUTAK-22.1-MDR is self-transmissible, whereas pUTAK-22.2-MDR and pUTAK-22.4-pO111 are most consistent with mobilization *in trans,* making use of the strain’s native helper plasmid background, while pUTAK-22.3-P1 represents a conjugation-deficient IncY phage-derived replicon. Genome annotation and comparative analyses further highlighted the structural diversity and mosaic composition of these plasmids, including P1-like phage remnants, virulence loci, and distinct antimicrobial resistance modules. Phenotypic profiling of the native wild-type plasmid complement and selected transconjugants in the recipient *E. coli* strain WG5 further showed that the presence of co-resident plasmids and redundant resistance determinants results in dosage-dependent streptomycin tolerance. Together, these findings expand our understanding of the mobility landscape, evolutionary dynamics, and resistance potential of STEC plasmids, while underscoring the importance of interpreting resistance phenotypes in the context of a strain’s natural plasmid composition.

## Introduction

Shiga toxin-producing *Escherichia coli* (STEC) are major zoonotic foodborne pathogens. STEC are responsible for illnesses ranging from mild diarrhea to hemorrhagic colitis (HC) and hemolytic uremic syndrome (HUS), with disease outcome shaped by the combined effects of chromosomal, phage-encoded, and plasmid-borne virulence determinants (<u>1–8</u>). Among these, serogroup O111 isolates comprise at least three pathovars: enteropathogenic *E. coli* (EPEC), STEC, including enterohemorrhagic *E. coli* (EHEC), and enteroaggregative *E. coli* (EAEC) (<u>9</u>, <u>10</u>). STEC of serotype O111:H8 represents a clinically significant lineage among the emerging “Big Six” non-O157 serogroups (<u>11</u>) linked to HUS (<u>12–19</u>). The STEC virulence hallmark is the production of potent toxins that function as translation inhibitors, encoded on lysogenic bacteriophages (<u>20–22</u>). Antibiotic exposure can induce the lytic cycle, leading to phage mobilization and ultimately to Stx expression (<u>23</u>, <u>24</u>). The Shiga toxin-converting bacteriophages (ΦStx) feature different combinations of *stx*-suballeles (<u>20</u>, <u>25</u>, <u>26</u>), among the two major toxin types, Stx1 and Stx2, differing in sequence, antigenicity, and associated disease severity (<u>27</u>). These phages introduce potent cytotoxins that confer a niche advantage during intestinal colonization in humans and in the bovine host by promoting the displacement of competing resident microbiota (<u>28</u>). Optimization of virulence and host adaptation has been the dominant selective pressure shaping the current STEC plasmidome (<u>29</u>). STEC plasmids are widely distributed among clinically important O157 and non-O157 serogroups, underscoring their central role in the ecology and evolution of virulence in STEC lineages (<u>8</u>, <u>30–32</u>). Among the plasmid-borne factors is enterohemolysin (Ehx), a pore-forming toxin that is a virulence hallmark of enterohemorrhagic *E. coli* (EHEC) pathotypes (<u>33</u>, <u>34</u>). The *ehxCABD* operon is highly conserved and syntenic across the major STEC serogroups, and in O111 is frequently associated with distinct IncFII or IncFIB backbones (<u>30</u>), underscoring its role as an evolutionarily stable virulence determinant (<u>35–37</u>). Other key plasmid genes include the serine protease autotransporter *espP* (<u>38</u>), the catalase–peroxidase *katP* (<u>39</u>), subtilase cytotoxin *subAB* (<u>40</u>), and adhesin or secretion-associated genes such as *toxB*, suggesting the prevalence of common virulence factors in plasmid lineages across the EHEC serogroups (<u>1</u>, <u>41</u>). Emerging antimicrobial-resistant (AR) and multidrug-resistant (MDR) strains within STEC populations pose challenges for treatment, surveillance, and public health control strategies (<u>42–45</u>). Multiple studies have reported MDR in outbreak-related strains (<u>45</u>). Consistent with this observation, AR and MDR O111 STEC isolates have been reported globally (<u>46–55</u>), though in general, MDR has been considered uncommon in STEC when compared to other *E. coli* pathovars (<u>8</u>, <u>42</u>, <u>45</u>, <u>56</u>). By definition, MDR-STEC plasmids encode resistance to three or more antibiotic classes, whereas AR-STEC plasmids carry resistance genes for fewer than three antibiotic classes (<u>57</u>). Plasmid-mediated horizontal gene transfer is a central mechanism driving the dissemination of antimicrobial resistance and virulence genes within *E. coli* and across bacterial species (<u>32</u>, <u>58–60</u>). Other mobile genetic elements (MGEs), such as bacteriophages (<u>22</u>, <u>61–64</u>) and transposons, also contribute to gene dissemination. Tn3/Tn21-family transposons are often associated with class 1 integrons (<u>65</u>) and insertion sequences (IS elements) such as IS*26* or ISCR*1*, which drive further rearrangement and mobilization of resistance loci (<u>66</u>). Antimicrobial resistance genes (ARGs) are frequently located on conjugative IncF and IncB/O/K/Z plasmid families (<u>41</u>, <u>64</u>, <u>67</u>) and documented in O111:H8 and non-motile O111 lineages (<u>29</u>, <u>68</u>). This organization promotes the mobilization and co-selection of virulence and antimicrobial resistance traits, contributing to MDR-STEC emergence and plasticity. Acquisition of IncB/O/K/Z and IncFII plasmids has been reported to impose little to no fitness cost in *E. coli* K-12, and some ARGs, such as *tetAR*, may even confer a fitness advantage under antimicrobial pressure (<u>29</u>, <u>69</u>). Understanding the prevalence, genetic context, and mobilization potential of MDR plasmids in STEC, as well as how antibiotic exposure influences ΦStx phage dynamics and Stx-production, is essential for advancing risk assessment, clinical management, and molecular epidemiology of this important pathogen group. In this study, we report the complete genome and plasmid repertoire of an MDR-STEC O111:H8 isolate, UTAK-22, which carries two MDR plasmids, a streptomycin-resistance/virulence plasmid, and a P1-like phage plasmid. We characterize the evolutionary and functional context of the plasmids, focusing on the conjugation machinery and genotype–phenotype correlations underlying antimicrobial resistance. We show how plasmid diversity enhances *E. coli* pathogenic potential and shapes its adaptive landscape. The intricate interplay between antimicrobial pressure, phage biology, and toxin regulation creates a dynamic environment in which resistance and virulence traits can converge. Antibiotic-induced stress responses may activate prophages, elevate Stx production, and exacerbate disease severity, underscoring the risks of inappropriate antimicrobial therapy (<u>70</u>, <u>71</u>).

## 2. Materials and Methods

### Bacterial strain analyzed in this study

For this study, the complete genome of the MDR-STEC of serotype O111:H8 strain UTAK-22, isolated from a veal carcass, was sequenced. Of the four carried plasmids, two were multidrug-resistant, one was streptomycin-resistant, and one was pan-susceptible. **Table S1** presents strain and genome-associated metadata.

### Genome sequencing, assembly, and annotation

Strains were cultured overnight at 37°C with shaking at 220 rpm in lysogeny broth (LB) (Thermo Fisher Scientific, Asheville, NC, USA). To maximize total genomic DNA (gDNA) yields, bacterial overnight cultures were diluted to an OD₆₀₀ of 0.03 in fresh LB medium and grown at 37°C with shaking at 220 rpm to mid-log phase (OD₆₀₀ ≈ 0.5). Total gDNA was extracted using the Monarch HMW DNA Extraction Kit (New England Biolabs, Ipswich, MA, USA). Genomes were sequenced to closure using Nanopore long-read technology (Oxford Nanopore, UK). Sequencing libraries were prepared using the Rapid Barcoding Kit (RBK-114) according to the manufacturer’s instructions and sequenced on the PromethION platform running a R10.4.1 Flow Cell (FLO-MIN114). Reads in the FASTQ format were imported into Galaxy v.22.05 (<u>72</u>). Default parameters were used for all software unless specified otherwise. FASTQ reads were quality-checked using FastQC (v.0.74+galaxy0) (http://www.bioinformatics.babraham.ac.uk/projects/fastqc) and assembled with Flye (v.2.9.3) (<u>73</u>). The resulting contigs were evaluated using QUAST (v.5.2.0+galaxy1). The chromosomal *dnaA* and plasmid *repA* genes were designated as the respective zero points of the closed molecules prior to annotation using the National Center for Biotechnology Information (NCBI) Prokaryotic Genome Annotation Pipeline (PGAP) (<u>74</u>). To estimate the number of respective plasmid copies, Oxford Nanopore PromethION reads were imported into Geneious Prime 2025.1.3 (GraphPad Software LLC; https://www.geneious.com). The closed UTAK-22 reference genome (GCA_041884315.1) was retrieved from the NCBI Assembly database and imported as a full GenBank record containing the chromosome and four plasmids. Reads were mapped to this combined reference using the Map to Reference workflow with Minimap2 v2.24 and the “PacBio/Oxford Nanopore” preset, with the default map-ont settings (https://github.com/lh3/minimap2). Geneious coverage statistics were used to calculate the mean depth for the chromosome and each plasmid. Plasmid copy number (PCN) was then calculated by normalizing plasmid mean coverage to the chromosomal mean coverage. Mean and median depths were nearly identical across replicons, confirming uniform coverage for each plasmid. For each plasmid, its copy number was calculated as the ratio of mean plasmid-to-chromosomal coverage. Because repetitive and conserved mobile-element regions may inflate coverage estimates, PCN values were interpreted as approximations.

### Phylogenomic analyses and MLST schemas

The genome, along with the sequence of K-12 substrain MG1655 (<u>75</u>, <u>76</u>), was imported into Ridom SeqSphere+ (v.8.3) (Ridom GmbH, Münster, Germany) for core genome (cg) and targeted Multilocus Sequence Typing (MLST) (<u>77–79</u>). The Sequence Type (ST) was determined according to the EnteroBase schema (<u>79</u>). Allele sequences for the Achtman scheme, targeting seven housekeeping genes (*adk*, *fumC*, *gyrB*, *icd*, *mdh*, *purA*, and *recA*), were accessed on the EnteroBase website (https://enterobase.warwick.ac.uk/species/ecoli/download_7_gene). A cgMLST schema was developed using the closed chromosome of the K-12 substrain MG1655 (<u>76</u>) as a seed, as previously described (<u>80</u>). Core and accessory MLST targets were identified according to the SeqSphere+ Target Definer’s inclusion/exclusion criteria. Allele information from the targeted seven-gene schema and the defined core genome genes of the panel strains was used to construct phylogenetic hypotheses using the minimum-spanning method with default settings (<u>81</u>, <u>82</u>).

### Pathogenome makeup and visualization

The chromosome and carried plasmids were compared and visualized using the Blast Ring Image Generator (BRIG; v.0.95) (<u>83</u>). To obtain complete *E. coli* chromosomal genomes, the NCBI Entrez Direct (EDirect) command-line toolkit (https://www.ncbi.nlm.nih.gov/books/NBK179288/) was used. A query was constructed to filter for assemblies labeled “latest” or “complete genome” and present in the RefSeq database at NCBI (https://www.ncbi.nlm.nih.gov/refseq/). Assembly records were retrieved using esearch/esummary, and metadata fields were parsed with xtract. To exclude plasmid-only assemblies, entries with "plasmid" in the FTP path were filtered out using grep. The remaining FTP paths were used to download compressed FASTA (*.fna.gz) files containing chromosomal sequences via wget. To identify the closest related genomes to the reference strain UTAK-22, the MinHash-based genome distance estimation tool Mash (v2.3) (<u>84</u>) was applied using default sketching parameters. Pairwise Mash distances were calculated between UTAK-22 and all available closed *E. coli* genomes. The 30 phylogenetically closest genomes with the lowest Mash distances were selected for further comparative analysis. The chromosomal serotype was determined *in silico* using ECTyper (v2.0.0+galaxy0) (https://github.com/phac-nml/ecoli_serotyping). Virulence genes and ARGs were cataloged using abricate (v1.4.0) (https://github.com/tseemann/abricate), querying the Virulence Factors of Pathogenic Bacteria (VFDB) (https://www.mgc.ac.cn/VFs/) (<u>85</u>), ResFinder (https://cge.cbs.dtu.dk/services/ResFinder/) (<u>86</u>) and Comprehensive Antibiotic Resistance Database (CARD) (https://card.mcmaster.ca/) (<u>87</u>). Ribosomal RNAs were detected with Basic Rapid Ribosomal RNA Predictor (barrnap v.1.10) (<u>88</u>). CRISPR/Cas systems were detected with CRISPRCasFinder (<u>89</u>) (v4.2.20) in Proksee (<u>90</u>) (v.1.1.0). Anti-phage systems were detected using DefenseFinder (<u>91</u>) (v.2.0.1+galaxy1). Loci of interest, such as ΦStx prophages, the Locus of Enterocyte Effacement (LEE) islands, and plasmids, were visualized and compared in pyGenomeViz-pgv-mauve (<u>92</u>) (https://github.com/moshi4/pyGenomeViz) using seaborn (<u>93</u>). Boundaries and locations of intact, partial, or remnant prophages were identified using PHASTEST (<u>94</u>), followed by manual curation of the ΦStx prophage and core genome borders by Mauve alignment of the genome to *E. coli* K-12 substrain MG1655 (GenBank accession U00096) (<u>75</u>) in Geneious Prime (v.2024.0.5). Direct repeats caused by phage integration were identified by BLASTn self-alignment and the Repeat Finder plugin (v1.0.1) in Geneious Prime. Replicase subtyping of the Stx-encoding prophages used the typing scheme introduced by (<u>95</u>). Replicases were typed using the EHEC phage replication unit (*eru*) schema (<u>96</u>, <u>97</u>). The subtypes of phage-borne *stx* were determined by BLASTn against a curated *stx*-suballele database, as previously described (<u>11</u>, <u>98–100</u>). A hierarchical classification approach was implemented to categorize phage-associated genes from the PHASTEST JSON output. All present phage-associated genes were further curated by incorporating processing type, name descriptors, and reported Gene Ontology (GO) terms to enable more in-depth functional classification. Genomic islands (GI) were detected with IslandViewer4 (<u>101–103</u>). LEE islands and boundaries were curated, guided by LEE1-LEE4 operon genes *espG and espF,* respectively, and visualized in pyGenomeViz-pgv-mauve (<u>92</u>) (https://github.com/moshi4/pyGenomeViz). The carried LEE intimin subtype was determined by BLASTn against a curated database of published *eae*-subtype sequences, as reported previously (<u>11</u>). Transposable elements and IS elements, along with inverted repeats, were cataloged through BLASTn (<u>104</u>) against the Transposon Central (TnCentral) database (<u>105</u>), comprising TnCentral+Integrall+ISFinder entries (<u>106</u>, <u>107</u>) and ISEScan (v.1.7.2.3+galaxy1) (<u>108</u>).

### Plasmid makeup and visualization

Plasmids were visualized and compared using the Blast Ring Image Generator (BRIG) (v.0.95) (<u>83</u>). Plasmid typing across taxonomic, mobility, and gene-inventory tiers was orchestrated with PlasmidTyper (<u>109</u>). Plasmid mobility and incompatibility groups were recorded using a local installation of Mobilome Typer (MOB-Typer) (v.3.1.9, installed via Bioconda) (<u>110</u>), and the per-plasmid reports were compiled by PlasmidTyper(<u>109</u>). This included the mobility features of DNA transfer, such as the type of relaxase (MOB) family and the Mating Pair Formation (MPF) system type I of the Type IV Secretion System (T4SS) (<u>110</u>). The following criteria were used: conjugative, presence of *oriT*, relaxase, type IV coupling protein (T4CP), and Type IV Secretion System (T4SS); non-mobilizable, absence of *oriT* or relaxase; and mobilizable, presence of *oriT* and relaxase, but absence of a complete T4SS. Plasmid Taxonomic Units (PTUs) were identified using COPLA, a plasmid taxonomic classifier, with default parameters (<u>111</u>). Using a Mash- and BLASTn-based search strategies, phylogenetically related plasmids were identified with the Plasmid Database software (PLSDB) (v.2024.05.31.v2) (<u>84</u>, <u>112</u>, <u>113</u>). Serotypes of the plasmid-associated chromosome molecules were determined *in silico* using ECTyper (https://github.com/phac-nml/ecoli_serotyping) (<u>114</u>). Virulence genes were cataloged with VFDB (<u>85</u>). Bacteriocins, ribosomally synthesized and post-translationally modified peptides (RiPPs), were recorded with BAGEL4 (http://bagel4.molgenrug.nl/) (<u>115</u>). Genomic islands were detected with IslandViewer4 (<u>101–103</u>). MGEs were identified with the mobile orthologous groups database software (mobileOG-db) (<u>116</u>). Integrons were detected and annotated using Integron Finder (https://github.com/gem-pasteur/Integron_Finder) (<u>117</u>). Transposable elements and IS elements were identified using ISEScan (<u>108</u>) and BLASTn (<u>104</u>) against the TnCentral database, comprising TnCentral+Integrall+ISFinder entries (<u>105–107</u>).

### Minimum Inhibitory Concentration (MIC) Assay

To determine the minimum inhibitory concentration (MIC) of selected antibiotics, a broth microdilution assay was performed in a 96-well plate format. LB broth was used as the growth medium for all conditions. Positive control wells contained bacteria in LB without antibiotics, while negative control wells contained LB broth alone to monitor for contamination. The plates were incubated in a plate reader at 37°C, and the OD₆₀₀ was measured every 10 min over 12 h. MIC values were defined as the lowest antibiotic concentration that completely inhibited visible bacterial growth. CLSI-recommended antibiotic concentration ranges were used, and overnight cultures were grown to mid-log phase (OD₆₀₀ ≈ 0.5) and then normalized to OD₆₀₀ of ≈ 0.01 (<u>118–120</u>).

### Plasmid Conjugation Assay

For the conjugation experiments, STEC UTAK-22 (TETᴿ STRᴿ GENᴿ NALˢ) served as the plasmid donor and *E. coli* WG5 Nalᴿ as the recipient strain (<u>121</u>). Overnight cultures were grown in LB at 37°C with aeration (200 rpm), then subcultured to an initial OD₆₀₀ of 0.05 in antibiotic-free LB and incubated until mid-log phase (OD₆₀₀ ≈ 0.5). Cultures were harvested (5,000 × g, 5 min), washed once in 1× PBS, and each pellet was resuspended in 1 mL LB. Donor and recipient suspensions were mixed at a 1:1 volumetric ratio to generate a 200 µL mating mixture per technical replicate (e.g., 100 µL donor and 100 µL recipient). The full 200 µL mixture was divided into four 50 µL mating spots on LB agar and incubated for 4 h at 37°C (<u>122</u>). Following mating, all four spots were collected together into 1× PBS using a sterile inoculation loop, pooled into a single tube, and vortexed to fully resuspend. Serial dilutions were prepared in 1× PBS from 100 to 10⁻⁶. For CFU quantification, 10 µL of each dilution was plated by vertical spotting onto LB agar supplemented with selective antibiotics (<u>123</u>). Nalidixic acid (50 µg/mL), along with the respective antibiotics, was used to enumerate transconjugants, while triple selection (TET+STR+GEN; 50 µg/mL) was used to quantify the donor population. Plates were incubated overnight at 37°C, and colonies were enumerated the next day. Plasmid identity of transconjugants was confirmed by colony PCR using plasmid-specific primers (**Table S2**). CFU/mL values were calculated by correcting for dilution and by scaling to the 10 µL spot volume. Conjugation efficiency was reported as transconjugants per donor using the following relationship: where NAL+Ab selects for transconjugants acquiring plasmid-encoded resistance, and TET+STR+GEN represents total donors recovered after mating. Conjugation frequencies were calculated as transconjugants per donor (T/D) for each mating. Values were log₁₀-transformed to account for the wide dynamic range of the data. For each plasmid, two technical replicate matings were performed per experiment and averaged to yield a single biological replicate. Log₁₀-transformed biological replicate values were analyzed using GraphPad Prism (v.10.5.0; GraphPad Software, San Diego, CA, USA). One-sample, two-tailed t-tests were performed for each plasmid. A *P* value ≤ 0.05 was considered statistically significant. STRᴿ CFU attributable to pUTAK-22.4-pO111 alone or in combination with pUTAK-22.1-MDR were estimated by subtracting CFU of pUTAK-22.1-MDR single transconjugants, and pUTAK-22.1-MDR/pUTAK-22.2-MDR co-transconjugants from total CFU recovered on STRᴿ plates; pUTAK-22.2-MDR alone yielded no colonies.

## Results and discussion

### Pathochromosome makeup and architecture

The genome of the Shiga toxin- and LEE-positive *E. coli* O111:H8 strain UTAK-22, isolated from a veal carcass, was sequenced to closure using Oxford Nanopore long-read technology, and consists of a chromosome and four carried plasmids that contribute antimicrobial resistance and virulence loci **(Table S1)**. The chromosome measures 5,303,756 bp in length with a GC content of 50.5%, typical for enteric STEC lineages **(Fig. 1)** (<u>11</u>, <u>124</u>). The plasmidome totals ∼420.9 kb and represents 7.35% of the 5.72-Mb total genome size. The four plasmids range in size from 66.9 kb to 129.3 kb, with GC contents from 47.5% to 53.5%, and exhibit diverse evolutionary plasmid backgrounds **(Fig. 2)**. To infer the phylogenomic context of this strain, the 30 most phylogenetically similar *E. coli* chromosomes based on Mash-distances were used to construct targeted MLST and cgMLST frameworks **(Fig. S1)**. The shared inventory comprises 4,160 loci, including 3,190 core and 970 accessory genes, which indicates a conserved *E. coli* backbone (<u>61</u>, <u>125–127</u>). High-resolution cgMLST typing clustered UTAK-22 within phylogroup O111:H8. The closest relative was identified as strain 2015C-3108 (O111:H8) of ST16, differing by 114 allelic changes **(Fig. S1)**, although UTAK-22 carries a novel *gyrB* allele not represented in the Achtman database, leaving its ST undetermined **(Table S3)**. The resulting phylogeny demonstrated relationships to other STEC serogroups, including three “Big Six” and other non-O157 emerging serogroups. Listed in order of cgMLST relatedness to UTAK-22: O111:H8 (sequence types ST16, ST294, and ST1792), O71:H11 (ST21), O26:H11 (ST21), O118:H16 (ST unknown), O45:H11 (ST29), O70:H11 (ST29), and O107/O117:H8 (ST2836) (<u>4</u>). Most isolates, representing H8, H11, and H16, were partitioned by H-antigen and ST, consistent with established patterns of *E. coli* evolution (<u>4</u>, <u>128</u>). A phylogenetic separation of O111 and O107/O117:H8 isolates was observed, defined by the positioning of H11 strains from serogroups O45 and O71. Previous studies placed O111:H8 (ST16), O26:H11 (ST21), and O118:H16 within the EHEC 2 clonal complex (<u>125</u>, <u>129</u>), a grouping recovered here. ST21 also included isolates of serotype O71:H11, and is the dominant sequence type among O26:H11 (<u>130</u>). The ST of O118:H16 is likewise unresolved, there because its seven called alleles, including a *purA* allele unique within the panel, form a combination not yet assigned an ST, though its allelic profile most closely resembles ST21 (<u>4</u>).

**FIG 1.**
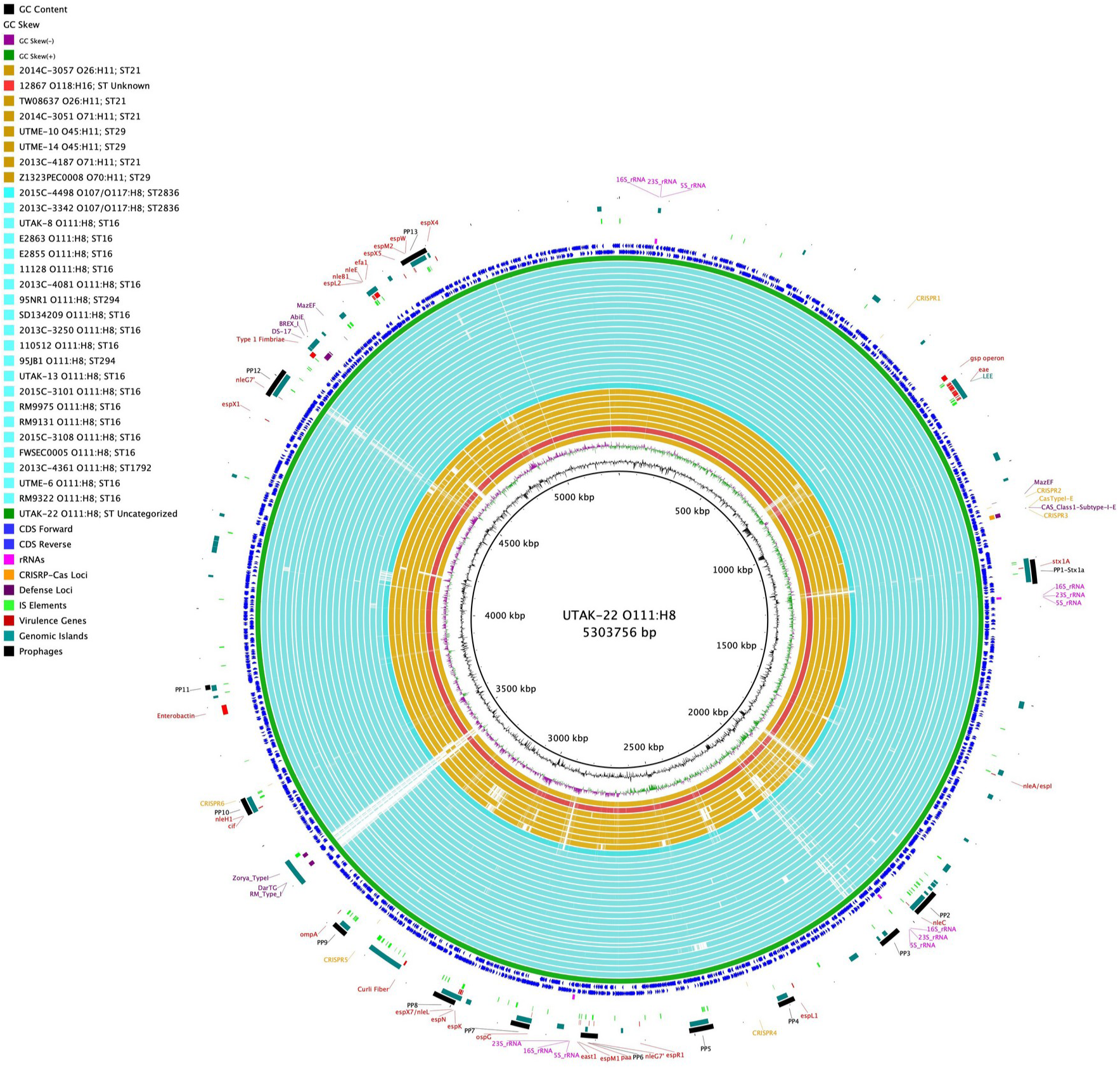
Comparison of the UTAK-22 chromosome with phylogenetically related *E. coli* genomes. BRIG comparison of the UTAK-22 chromosome with the 30 most closely related representative complete *E. coli* genomes identified by Mash-based similarity analysis. Query chromosomes are color-coded by serotype/lineage as indicated in the legend, and their order reflects inferred phylogenomic relatedness. Coding sequences on the positive and negative strands, GC content/skew, virulence-associated loci, genomic islands, prophages, and other loci of interest are highlighted as indicated in the legend.

**FIG 2.**
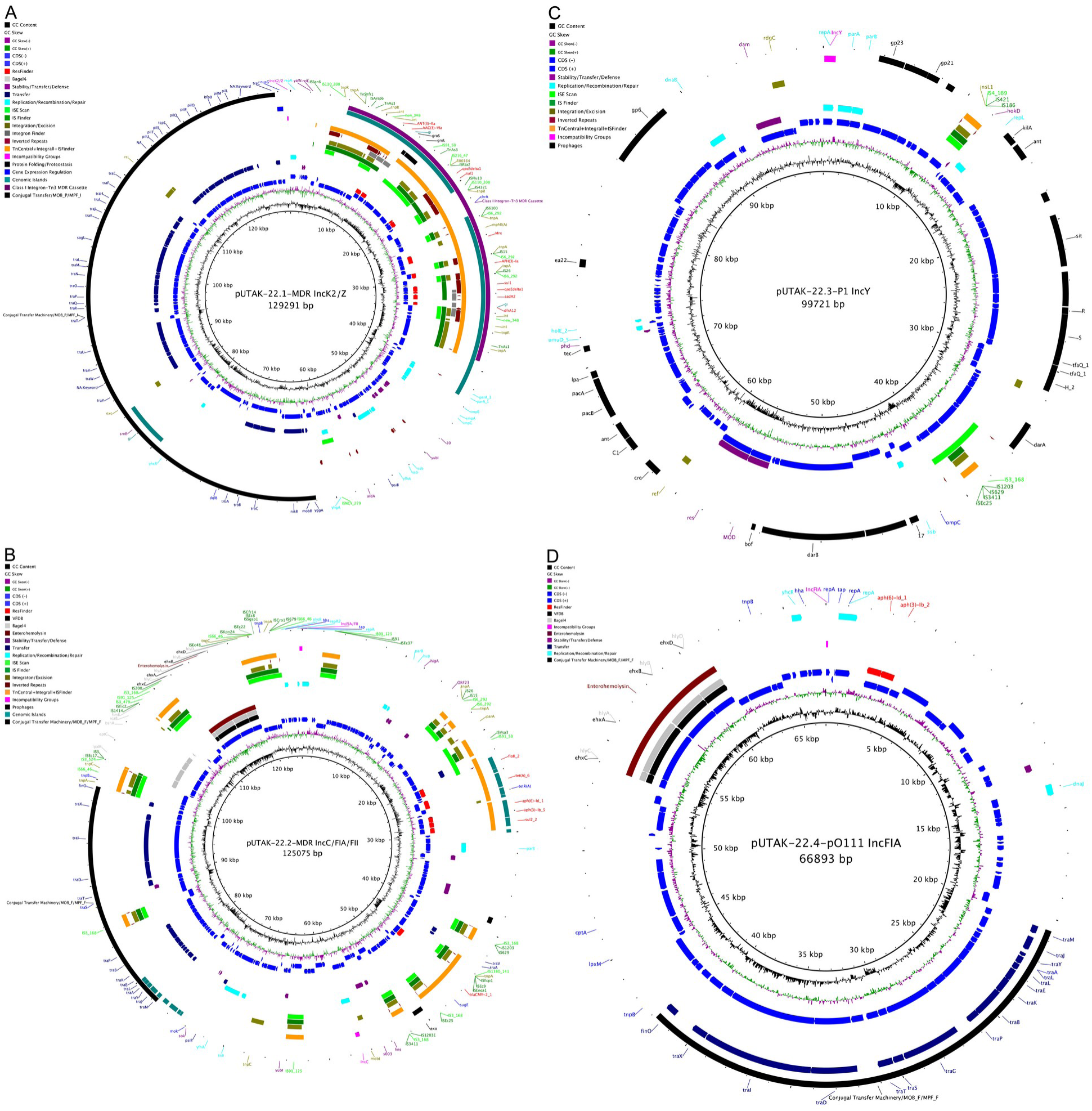
Plasmid architecture of the UTAK-22 plasmidome. BRIG visualization of the four plasmids carried by STEC strain UTAK-22. Coding sequences are shown on the positive and negative strands, and loci associated with antimicrobial resistance, virulence, replication/stability, insertion sequences, prophage-related functions, and conjugative transfer are highlighted as indicated in the legend. **(A)** pUTAK-22.1-MDR is a 129,291-bp IncK2/Z multidrug-resistant plasmid with a GC content of 53.5%. **(B)** pUTAK-22.2-MDR is a 125,075-bp hybrid IncC/FIA/FII multidrug-resistant plasmid with a GC content of 51.5%. **(C)** pUTAK-22.3-P1 is a 99,721-bp IncY P1-like phage-plasmid with a GC content of 47.5%. **(D)** pUTAK-22.4-pO111 is a 66,893-bp IncFIA O111-associated virulence plasmid with a GC content of 49.0%. Together, these maps illustrate the mosaic plasmid architecture of UTAK-22, including multidrug-resistance regions, virulence-associated cargo, phage-plasmid functions, and distinct transfer-related modules.

### Chromosome-embedded virulence determinants

Prophages and other identified pathogenicity-associated islands (PAI) make up 20.8% of the chromosomal coding capacity, a proportion similar to that observed in other STEC lineages (<u>11</u>, <u>61</u>) **(Fig. 1)**.

### Cryptic ΦStx1a prophage

A single cryptic ΦStx1a prophage was identified in UTAK-22 at the *ssrA* locus, a documented phage insertion site in STEC O111 (<u>131</u>). The prophage boundaries are marked by perfect 10-bp direct repeats **(Fig. 3A, Table S4)**. Bacteriophages commonly target conserved chromosomal loci and undergo acquisition, loss, and microevolution, processes that collectively shape gene content and pathogenicity traits in STEC (<u>22</u>, <u>132</u>, <u>133</u>). BLASTn comparisons indicated that ΦStx1a was most similar to the cryptic phages O111:NM and O111:H8, CP-1639, with ≥ 99% identity and 100% coverage (<u>131</u>). Although ΦStx bacteriophages are classically MGEs, cryptic ΦStx1 prophages harbor defects that may compromise autonomous propagation due to extensive gene decay and IS element disruption. However, experimental evidence indicates that inter-prophage interactions may restore the capacity of ΦStx1 prophages for excision, packaging, and horizontal transfer (<u>134</u>). Several loci are pseudogenized, including the tail assembly protein T, the tail-tube terminator GpU, and the C-terminal antirepressor KilAC. Three IS629 elements are present, one of which disrupts a tail fiber gene. The prophage also lacks multiple recombination components, including integrase and excisionase, as well as capsid genes. Its EHEC phage replication unit corresponded to Eru subtype 3 and contained a distinct replication module that replaced the canonical O/P region (<u>96</u>, <u>97</u>). This organization has been reported in *stx2*+ O111 and O157:H7 isolates (<u>96</u>). In ΦStx2a phage contexts, α-replicase variants have been associated with elevated Stx2 production relative to β- or δ-type replicases, but lower than that observed with γ-type (<u>95</u>).

**FIG 3.**
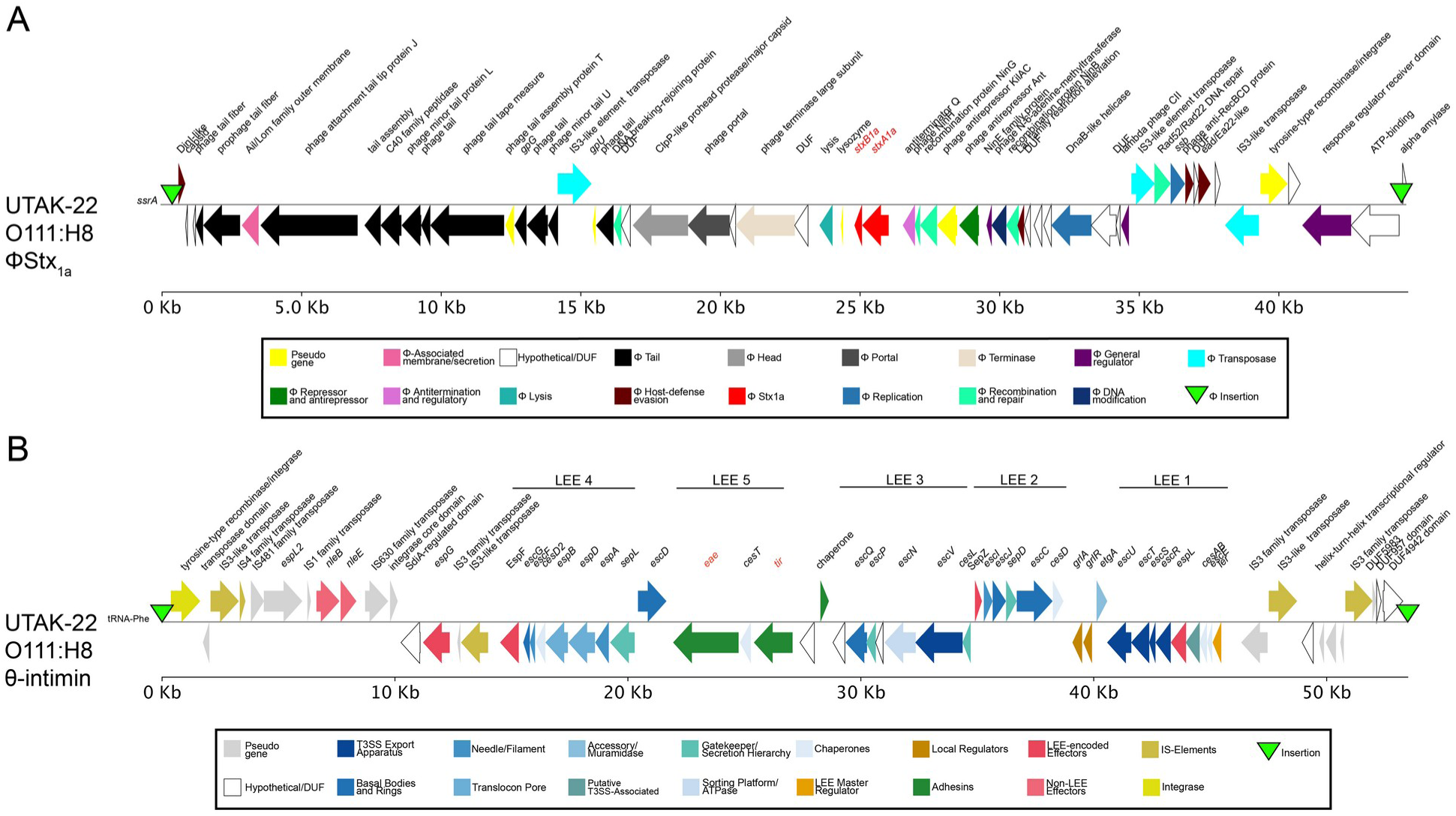
Chromosomal virulence determinants of UTAK-22. Comparative gene maps of the two major chromosomal virulence loci in UTAK-22. **(A)** The cryptic ΦStx1a prophage is approximately 43.0 kb (42,977 bp) in size and inserted at the tmRNA ssrA locus. **(B)** The approximately 52.9-kb LEE island is integrated at the tRNA-Phe locus, retains the canonical LEE1–LEE5 organization, and carries the θ-intimin subtype. Functional annotations of key phage, secretion, regulatory, and virulence-associated genes are highlighted as indicated in the legend.

### Locus of Enterocyte Effacement

The LEE PAI is prevalent in several STEC serotypes and is associated with the formation of characteristic attaching and effacing (A/E) lesions (<u>135–137</u>); and LEE carriage has been associated with HUS (<u>138</u>, <u>139</u>). In UTAK-22, the island features the θ-intimin subtype and is integrated at the tRNA-Phe locus **(Fig. 3B)**, a well-documented insertion site in *E. coli* (<u>140–142</u>), particularly in serogroup O111 (<u>143</u>). The island boundaries are defined by perfect 17 bp direct repeats **(Table S4)**. This island is organized into five polycistronic operons (LEE1 to 5), encoding type III secretion system (T3SS) regulators and effectors, together with non-LEE-encoded (Nle) effectors, transporters, and regulatory elements promoting intracellular survival, adhesion, immune modulation, and enterotoxin production (<u>139</u>, <u>144</u>, <u>145</u>).

### Plasmidome and associated virulence and resistance traits

The UTAK-22 plasmidome represents 7.35% of the total genome size and integrates 23 virulence and antimicrobial resistance determinants across four plasmids **(Table S5)**. The two largest replicons, pUTAK-22.1-MDR (129,291 bp; 53.5% GC) and pUTAK-22.2-MDR (125,075 bp; 51.5% GC), conferred MDR, each encoding resistance to at least three distinct antibiotic classes (<u>146</u>). The phage-like plasmid pUTAK-22.3-P1 (99,721 bp; 47.5% GC) did not encode ARGs, whereas the smallest serogroup-characteristic virulence plasmid, pUTAK-22.4-pO111 (66,893 bp; 49.0% GC), carried a single streptomycin (*str*) resistance locus (<u>143</u>). Coverage analysis normalized to chromosomal depth (541×) indicated that all four pUTAK-22 plasmids are maintained at low relative copy number. Estimated plasmid-to-chromosome coverage ratios were calculated as 0.61 for pUTAK-22.1-MDR, 1.41 for pUTAK-22.2-MDR, 1.88 for pUTAK-22.3-P1, and 1.48 for pUTAK-22.4-pO111. These values were consistent with their respective IncK2/Z, IncF, and IncY incompatibility-group classifications, which are common among *Enterobacteriaceae* and support low-copy plasmid status **(Table S5)** (<u>147–151</u>).

### Plasmid pUTAK-22.1-MDR

The largest plasmid, pUTAK-22.1-MDR (129,291 bp; 53.5% GC) belonged to the IncI-complex (IncB/O/K/Z) and Plasmid Taxonomic Unit (PTU) PTU-B/O/K/Z **(Fig. 2A)**. Plasmids in this clade typically have a narrow host range, largely restricted to *Enterobacteriaceae*, and are characterized as low- to medium-copy plasmids that are known to accumulate mobile genetic resistance elements (<u>111</u>, <u>152</u>, <u>153</u>). Consistent with this plasmid family, a relative copy number of 0.61 compared to the chromosome was calculated. The plasmid backbone includes conserved inheritance and maintenance modules, such as a ParB/RepB/Spo0J-family partitioning protein and multiple toxin–antitoxin systems that support stable maintenance. These systems comprised a Hok/Gef-family Type I toxin, a DinQ-like toxin dqlB, and a type II Phd/YefM–RelE/ParE module positioned adjacent to the replication region. Together, these systems form a multilayered stability network characteristic of large *Enterobacteriaceae* plasmids (<u>154</u>). This group is widely distributed among enteric bacteria and is often implicated in the dissemination of antimicrobial resistance determinants (<u>32</u>). A prominent conjugation and mobilization region spanning roughly 35 kb was identified, consisting of *tra/trb* genes assigned to the MOB_P and MPF_I systems, consistent with the transfer architecture of I-complex plasmids, such as IncK2/Z (<u>153</u>, <u>155</u>). Plasmids with MPF_I systems exhibit a narrow host range, primarily within the genera *Escherichia* and *Salmonella* (<u>32</u>, <u>156</u>). Five resistance genes confer resistance to three antibiotic classes: aminoglycosides, sulfonamides, and trimethoprim. Among these, the *sul1* and *aadA2* genes are present in duplicate, which suggested potential dosage effects.

### Modular class 1 integron–Tn3 MDR-cassette architecture in plasmid pUTAK-22.1-MDR

This plasmid carries a composite MDR region conferring resistance to aminoglycosides, sulfonamides, and trimethoprim **(Fig. 4)**. The aminoglycoside N-acetyltransferase *aac*(*3*)*-VIa* mediates resistance to gentamicin, whereas the aminoglycoside phosphotransferase *aph(3′)-Ia* protects against kanamycin and neomycin. A third enzyme, adenyltransferase *aadA2*, confers additional resistance to streptomycin and spectinomycin, thereby broadening the spectrum of aminoglycoside resistance. The plasmid further harbors a macrolide-resistance cluster, *mphR(A)–mrx(A)–mph(A)*, mobilized via IS26-associated transposition and commonly inserted near the 3′-conserved segment of class 1 integrons (<u>157</u>). In pUTAK-22.1-MDR, *mph(A)* is truncated, consistent with IS26-mediated recombination at this locus. The sulfonamide-resistance gene *sul1* is present in two copies, one within each class 1 integron, which mediates resistance to sulfisoxazole (SFX). Finally, the dihydrofolate reductase gene *dfrA12* confers trimethoprim (TMP) resistance. The co-occurrence of *sul1* and *dfrA12* supports the presence of class 1 integron structures, frequently embedded within Tn21 derivatives or composite resistance islands (<u>65</u>, <u>66</u>, <u>158</u>). Class 1 integrons are recognized as major genetic elements facilitating the acquisition and dissemination of antimicrobial resistance (<u>158</u>, <u>159</u>). ARGs were found to be embedded in two tandem class 1 integrons, each containing *intI1*, the *attI1* primary recombination site, *Pc/Pi* promoters, *attC* recombination sites, and the 3′-conserved segment featuring *qacEΔ1* and *sul1* as characteristic hallmarks (<u>66</u>, <u>160–162</u>) **(Fig. 4)**.

**FIG 4.**
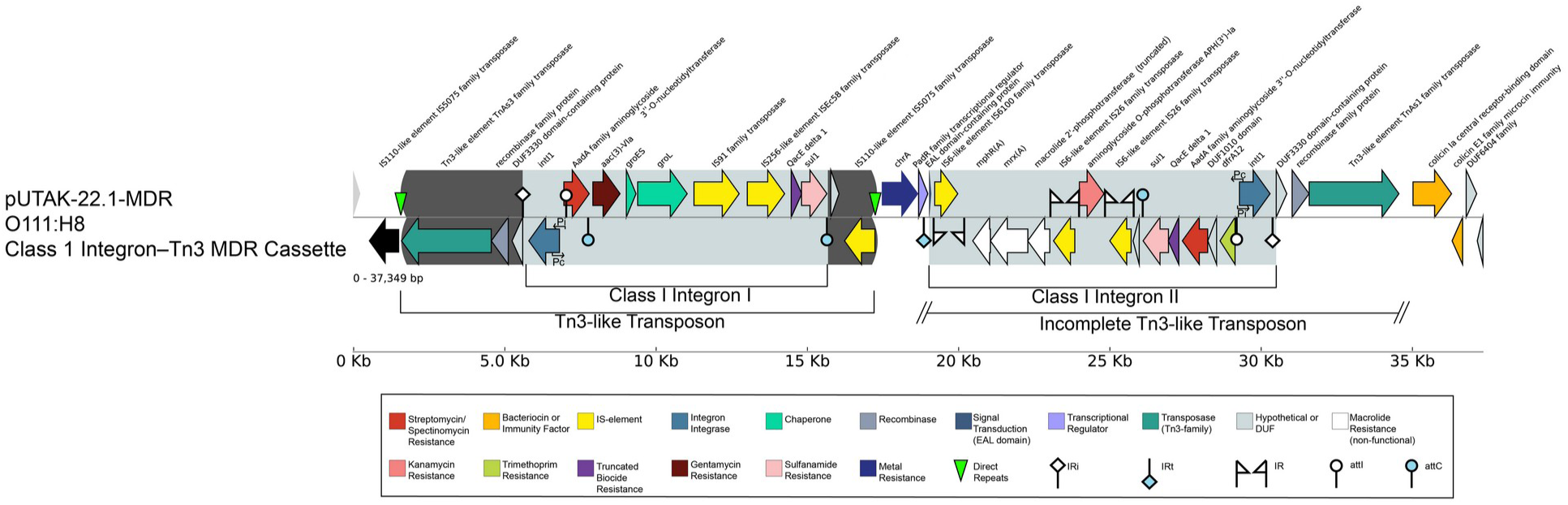
Genetic organization of the class 1 integron–Tn3 composite multidrug-resistance cassette in pUTAK-22.1-MDR. Gene map of the approximately 37-kb multidrug-resistance region of pUTAK-22.1-MDR. The cassette comprises two Tn3-like transposon-associated regions, each carrying a class 1 integron, embedded within an IS-rich segment containing IS26, IS6100, IS5075, and IS91-like elements. Together, the tandem integrons encode resistance determinants associated with streptomycin, gentamicin, trimethoprim, sulfonamides, quaternary ammonium compounds, kanamycin/neomycin, and the inducible macrolide-resistance operon *mphR(A)–mrx(A)*–*mph(A)*, the latter truncated by IS26. Both integrons retain canonical class 1 features, including *intI1*, the *attI1* primary recombination site, *attC* sites delimiting gene cassettes, and the 3′ conserved segment (*qacEΔ1*-*sul1*).

### Phylogenomic position of pUTAK-22.1-MDR

Plasmid pUTAK-22.1-MDR is classified as an IncK2/Z replicon (PTU-B/O/K/Z), a tightly clustered clade of O111:H8 EHEC plasmids with over 99% nucleotide identity across 90% of the backbone, including replication, stability, and transfer regions, that show potential introgression from other *Enterobacteriaceae* plasmids **(Fig. 5A)** (<u>111</u>, <u>153</u>). Nucleotide-level alignment revealed significant localized homology to the *repA-parAB-hok/pem* replication and stability module and *tra/trb* transfer region in a *Salmonella enterica* serovar. Two *K. pneumoniae* strains further displayed homology to the class I integron multidrug-resistance island of pUTAK-22.1-MDR, encoding a subset of the UTAK-22 AR cassette loci, including *ant(3″)-Ia*, *aac*(*3*)*-VIa*, *aph(3′)-Ia*, *aadA2*, *sul1*, and *dfrA12*. Taken together, our observations suggested that pUTAK-22.1-MDR is a chimeric mobilizable hub of AR dissemination. The plasmid likely descended from the O111:H8 EHEC lineage, whereas the resistance cassettes may have been horizontally acquired through successive interspecies exchanges with other *Enterobacteriaceae* pathogens. This interpretation was supported by the presence of similar hybrid plasmids, in which IncB/O/K/Z backbones acquired class 1 integrons from *Enterobacteriaceae* reservoirs (<u>32</u>, <u>59</u>).

**FIG 5.**
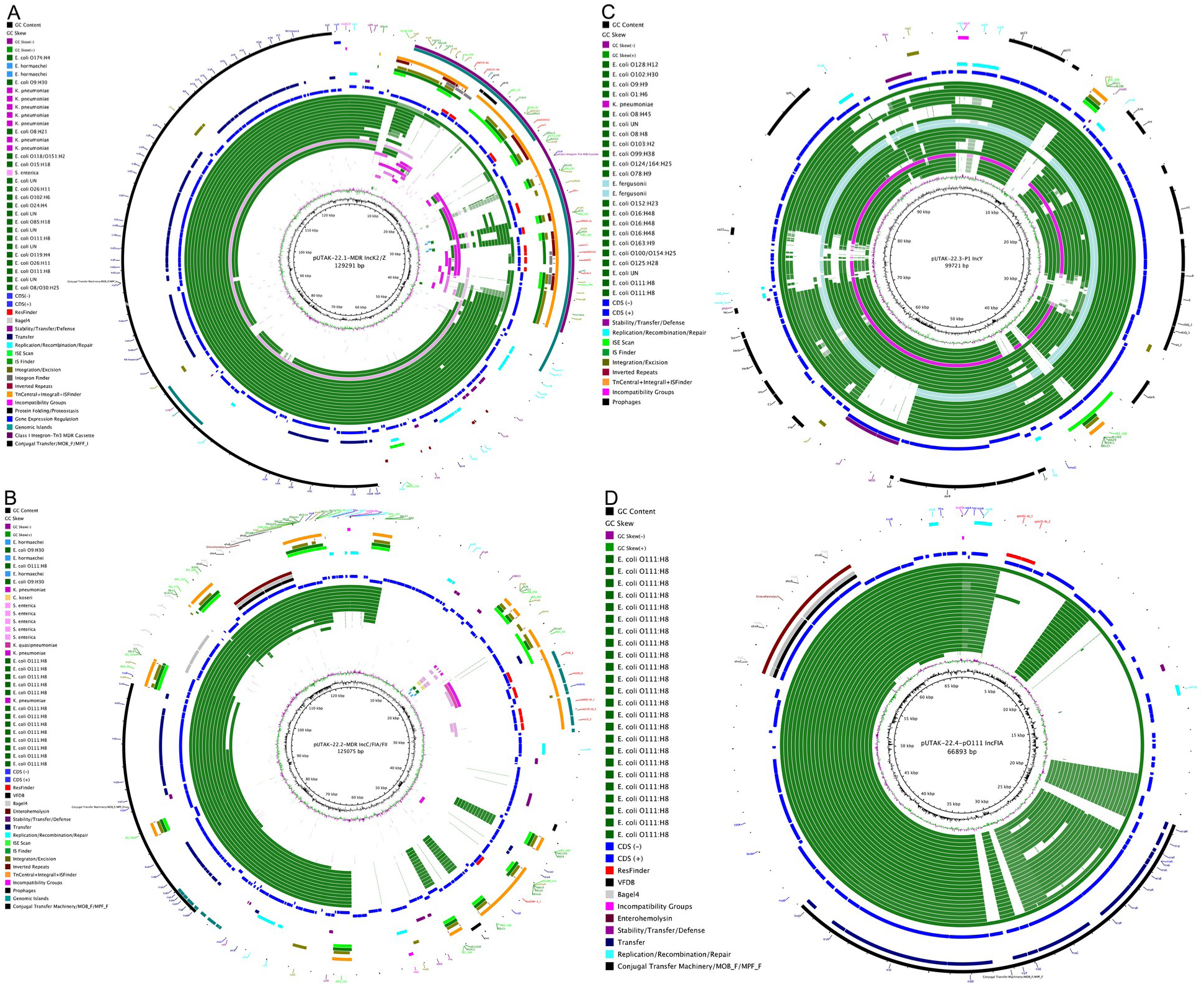
Comparative genomics of the UTAK-22 plasmidome and related plasmids. BRIG comparisons of the four UTAK-22 plasmids with related plasmids identified by Mash/PLSDB-based analyses. Query plasmids are color-coded by *E. coli* serotype or by non-*E. coli* species and are arranged according to inferred relatedness. Coding sequences and key loci associated with antimicrobial resistance, virulence, replication/stability, insertion sequences, prophage functions, and conjugative transfer are annotated as indicated in the legend. **(A)** pUTAK-22.1-MDR shares a conserved *Enterobacteriaceae* backbone, whereas its class 1 integron–Tn3 multidrug-resistance region shows an incoherent distribution across related plasmids. **(B)** pUTAK-22.2-MDR is most similar to related *E. coli* plasmids but also contains IncC-associated regions characteristic of broader *Enterobacteriaceae* exchange. **(C)** pUTAK-22.3-P1 groups with related IncY/P1-like phage-plasmids, primarily from diverse *E. coli* lineages, with more limited conservation in other *Enterobacteriaceae*. **(D)** pUTAK-22.4-pO111 is closely related to conserved O111:H8-lineage virulence plasmids but carries an acquired streptomycin-resistance locus.

### Hemolytic and conjugal virulence and antimicrobial resistance plasmids pUTAK-22.2-MDR and pUTAK-22.4-pO111

The 125,075 bp pUTAK-22.2-MDR plasmid (51.5% GC) and the smaller 66,893 bp pUTAK-22.4-pO111 plasmid (49.0% GC) were identified as F-family plasmids of the IncC/FIA/FII and IncFIA subtypes, respectively. The latter belonged to PTU-E41, whereas the PTU of pUTAK-22.2-MDR is undetermined (<u>148</u>, <u>163</u>). Comparative sequence inspection revealed that both plasmids share high nucleotide identity and conserved structural organization across replication and conjugation regions. The ∼20-kb transfer locus comprises a canonical IncF-type MOB_F/MPF_F cassette typical of self-transmissible F-type conjugative *E. coli* plasmids (<u>148</u>). pUTAK-22.4-pO111 encodes the canonical MOB_F/MPF_F cassettes characteristic of O111-associated virulence plasmids (<u>110</u>, <u>143</u>, <u>164</u>) **(Table S5)**. Both plasmids are maintained at low relative copy number, consistent with their Inc types and estimated copy numbers (<u>1</u>, <u>30</u>) (Table S5). An EHEC-type α-hemolysin (*ehx*) operon is present on both plasmids (<u>33</u>, <u>35–37</u>), along with a chromosomal enterobactin *ent-fep-fes* siderophore system (<u>165–167</u>). The presence of *ehxCABD* on both plasmids suggests possible gene-dosage effects enhancing hemolytic capacity (<u>1</u>, <u>2</u>). The perfect sequence conservation of the *ehxCABD* operons on both plasmids suggests an inter-plasmid exchange event rather than independent acquisition (<u>30</u>, <u>168</u>). In both plasmids, the operon is embedded within a composite IS-*hly*-IS module, a structure characteristic of transposable virulence cassettes mobilized through IS-mediated recombination (<u>65</u>, <u>169–171</u>). Considering that the *ehx* locus is an integral part of pO111 (<u>143</u>), it is possible that this locus was introduced by lateral exchange to pUTAK-22.2-MDR, similar to dissemination of an arsenic-resistance transposon among plague plasmids (<u>172</u>). Despite overall similarity, pUTAK-22.2-MDR carries an IS1203 insertion that disrupts the C-terminal domain of *traG* **(Fig. 2B, Fig. S2)** (<u>173</u>). This insertion pattern aligns with models of virulence-plasmid exchange involving pO111-like backbones (<u>143</u>). Both plasmids also contain accessory modules, including toxin–antitoxin systems, transposases, and metabolic genes, alongside homology to IncF-type plasmids circulating among STEC and related *Enterobacteriaceae*. While structurally similar, they differed in replicon composition, carrying complementary IncF replicons **(Table S5)** (<u>174</u>). This architecture is consistent with documented modular evolution and recombination events common among plasmids of *E. coli* O111 and O26 lineages (<u>30</u>, <u>175</u>). pUTAK-22.2-MDR additionally harbors an IncC replicase, *tra* remnants, and MDR loci, features consistent with mosaic IncC plasmids that circulate widely among *Enterobacteriaceae* species (<u>176–179</u>) **(Fig. 2B**, **Fig. 5B)**.

### Serogroup-specific plasmid pUTAK-22.4-pO111

Plasmid pUTAK-22.4-pO111 (66,893 bp; 49.0% GC) belonged to a distinct clade of virulence plasmids, prevalent in O111:H8/non-motile (H-) strains (<u>143</u>) **(Fig. 5D)**, and was classified within PTU-E41 (<u>1</u>, <u>30</u>, <u>111</u>). This group shares a conserved virulence-plasmid replication module, with similar size ranges (∼64-68 kb) and GC contents (∼49%) (<u>143</u>, <u>148</u>, <u>179</u>). The plasmid encodes a ∼20 kb transfer region containing the canonical IncFIA MOB_F/MPF_F cassette. This plasmid is widely distributed and conserved among O111 strains and enhances host colonization, toxin secretion, and immune evasion (<u>59</u>, <u>180</u>). pUTAK-22.4-pO111 is maintained at low-copy number and represents an IncFIA-type *E. coli* virulence replicon (<u>148</u>). Its replication and conjugation backbone is shared with pUTAK-22.2-MDR **(Fig. S2)**. Additionally, pUTAK-22.4-pO111 carries an aminoglycoside-resistance cassette composed of *aph*(*6*)*-Id* and *aph(3′)-Ib*, which confers streptomycin resistance (<u>181</u>). Although the pO111 plasmid backbone is highly conserved, the UTAK-22 variant had acquired an additional streptomycin-resistance locus also detected on pUTAK-22.2-MDR. This configuration is typical of a classical *strA-strB Enterobacteriaceae* plasmid cassette flanked by IS91 and Tn3-family transposases (<u>32</u>, <u>65</u>). The myristoyltransferase enzyme (*lpxM*) strengthens the outer membrane, thereby enhancing antimicrobial resistance, colonization, and persistence (<u>182</u>, <u>183</u>). Further, a phosphoethanolamine transferase (*cptA*) contributes to outer-membrane modification and overall fitness (<u>1</u>, <u>184</u>).

### Phage-like plasmid pUTAK-22.3-P1

The 99.7-kb plasmid, pUTAK-22.3-P1 (47.5% GC), was classified within PTU-Y and corresponded to the family of P1-like phage-plasmids **(Fig. 2C)** (<u>185</u>). These low-copy extrachromosomal elements are prevalent in *E. coli* and other *Enterobacteriaceae* (<u>163</u>). pUTAK-22.3-P1 clusters tightly (identity ≥ 99%, coverage ≥ 95%) with other P1 plasmids **(Fig. 5C)**. Highest nucleotide identities were observed among *E. coli* plasmids from diverse serotypes (e.g., O111:H8, O125:H28, O100/O154:H25, O163:H9, O16:H48, and O152:H23) and among avian pathogenic *E. coli* (APEC) plasmids, which share both P1-like phage modules and plasmid replication/maintenance loci. These similarities indicate overall backbone-level homology within *E. coli*. More distantly related matches to *E. fergusonii* and *Klebsiella pneumoniae* plasmids are limited to core conserved P1-like phage modules. A defining feature is the presence of two replicons: the archetypal type IncY replicon and a serogroup-specific pO111 replicon (<u>186</u>, <u>187</u>). Although P1-like plasmids can acquire AR or virulence genes through IS-mediated recombination, pUTAK-22.3-P1 did not carry any such determinants (<u>64</u>). Most of its gene inventory is phage-derived, including loci for phage morphogenesis, immunity, lysis, and packaging; features characteristic of P1-like phage plasmids (<u>164</u>, <u>188</u>). P1 plasmids have been widely associated with STEC and other enteropathogenic lineages, where they contribute to genomic plasticity, horizontal gene flux, and long-term maintenance (<u>143</u>, <u>189–191</u>). Consistent with its phage-plasmid hybrid origin, the replicon displays a relatively low GC content (47.5%) and carries plasmid maintenance determinants such as replication initiation *repA* and a *parA-parB* partitioning system, along with type I and II toxin-antitoxin modules (<u>188</u>). Several additional genes, including *traT* and *iss,* confer complement resistance and serum survival and are also prevalent in avian and human pathogenic *E. coli* (<u>30</u>, <u>192</u>, <u>193</u>). Housekeeping and stress-response genes, including *ompC*, *dam*, *rdgC*, and *dnaB*, contribute to membrane integrity, DNA replication stability, and oxidative-stress management (<u>194–197</u>).

### Conjugation efficiency of carried UTAK-22 plasmids

Conjugation assays were used to evaluate plasmid transfer capability and efficiency using nalidixic acid plus antibiotic-selective readouts in *E. coli* WG5 recipients (**Fig. 6**). *In silico* profiling predicted that pUTAK-22.1-MDR, pUTAK-22.2-MDR, and pUTAK-22.4-pO111 are potentially mobile, whereas pUTAK-22.3-P1 lacks identifiable transfer genes (**Table S5**). Experimental matings confirmed that pUTAK-22.1-MDR (IncK2/Z, MOB_P, MPF_I) is conjugative. The IncF-type plasmids pUTAK-22.2-MDR and pUTAK-22.4-pO111 belong to a highly prevalent conjugative plasmid group in *E. coli*, particularly among STEC/EHEC and EPEC pathotypes (<u>30</u>, <u>148</u>). These two plasmids show conserved synteny and high nucleotide identity across their *tra/trb* operons and associated regulatory loci (*traJ-finO-finP*) **(Fig. S2)**. Comparative genomics, however, revealed that their shared F-like transfer region is incomplete, comprising only ∼42% of the canonical region. These plasmids retained genes required for relaxosome formation, coupling, core T4SS assembly, regulation, surface exclusion, and post-transfer processing, but lacked the pilus-biogenesis genes (*traV/C/W/U/F*) required for autonomous F-like conjugation. Consistent with this architecture, pUTAK-22.2-MDR was non-conjugative but was recovered as a co-mobilized plasmid in the presence of pUTAK-22.1-MDR. The observed transconjugant frequencies fell within the typical range reported for self-transmissible and mobilizable plasmids in EHEC and related pathogens (<u>30</u>, <u>147</u>, <u>149</u>) **(Fig. 6, Table S6)**. Direct assessment of pUTAK-22.4-pO111 transfer was impeded by the shared STRᴿ markers across the plasmid set. Selection on STRᴿ, followed by pooling of transconjugants and multiplex PCR screening indicated the presence of pUTAK-22.1-MDR along with pUTAK-22.2-MDR and pUTAK-22.4-pO111. These findings demonstrated that pUTAK-22.4-pO111 is transferable in this plasmid context (data not shown), although its incomplete transfer region suggests mobilization *in trans* rather than conjugation. This model aligns with previous observations that non-conjugative plasmids may exploit conjugation machinery supplied by co-resident plasmids (<u>58</u>, <u>185</u>, <u>186</u>).

**FIG 6.**
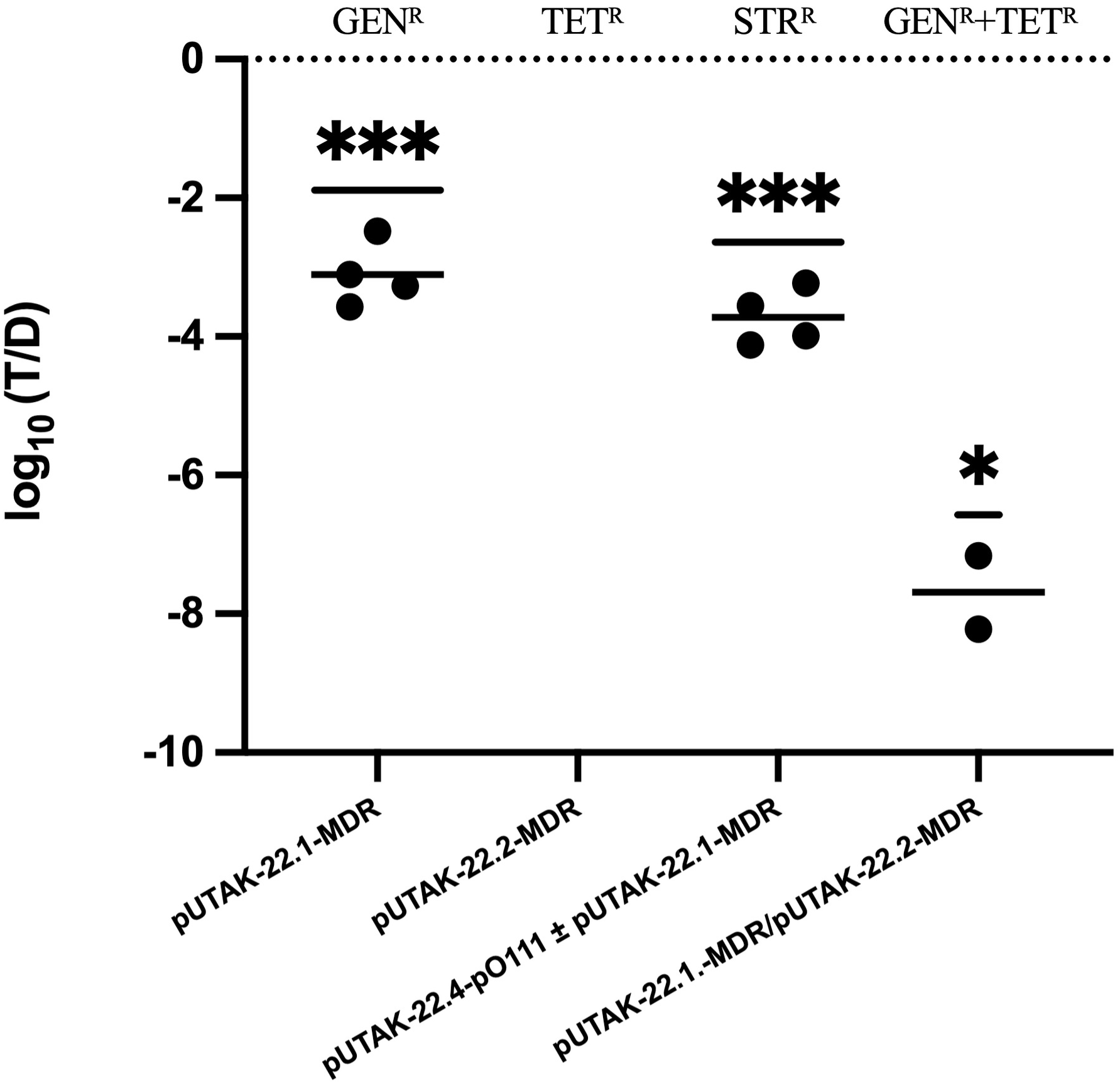
Conjugation frequencies of UTAK-22 plasmids. Log₁₀-transformed conjugation frequencies, expressed as transconjugants per donor (T/D), are shown for pUTAK-22.1-MDR, pUTAK-22.4-pO111, and the pUTAK-22.1-MDR/pUTAK-22.2-MDR co-transfer condition. Each point represents an independent biological replicate, where each biological value is the mean of two technical replicate matings, and horizontal lines indicate the mean. The dotted line at 0 corresponds to a theoretical conjugation frequency of T/D = 1. pUTAK-22.2-MDR alone yielded no detectable colonies under the assay conditions. For each condition with recovered transconjugants, log₁₀(T/D) values were compared with a theoretical mean of 0 using one-sample, two-tailed t-tests in GraphPad Prism 10 (\*\*\**P* ≤ 0.001; \**P* ≤ 0.05).

Although pUTAK-22.3-P1 lacks an antibiotic resistance marker that would permit direct selection and enumeration of transconjugants, multiplex PCR analysis of pooled streptomycin-resistant transconjugants detected pUTAK-22.3-P1 within the transconjugant population. Thus, transfer of pUTAK-22.3-P1 was detected qualitatively, but an independent transfer frequency could not be determined. This finding is notable given its predicted non-mobilizable status and is consistent with reports that P1-like plasmids can be mobilized through fusion with conjugative plasmid backbones (<u>198</u>).

### The antimicrobial resistance gene inventory of UTAK-22

Strain UTAK-22 harbors three plasmids encoding ARG loci, two of which are classified as MDR plasmids **(Fig. 2, Table S5)**. The 129-kb MDR plasmid pUTAK-22.1-MDR encodes five distinct resistance genes, present in single or double copies, that collectively confer protection against three antibiotic classes **(Fig. 2A)**. These loci includes aminoglycoside-modifying enzymes conferring resistance to gentamicin (GEN; 1× *aac*(*3*)*-VIa*), kanamycin (KAN; 1× *aph(3′)-Ia*), and streptomycin (STR; 2× *aadA2*), together with the sulfonamide resistance gene (2× *sul1*) and the trimethoprim (TMP) resistance determinant (1× *dfrA12*), encoding a dihydrofolate reductase (<u>199–203</u>). This plasmid also encodes the macrolide-resistance operon *mphR(A)–mrx(A)–mphA* (<u>204</u>). However, *mphA* carried a frameshift mutation that caused premature truncation at amino acid residues Δ113-243, disrupting the N-terminal ATP-binding region and likely abolishing function **(Fig. 4)** (<u>205</u>). Despite this, UTAK-22 remains erythromycin (ERY) resistant **(Fig. S3)**. CARD analysis identified an intact chromosomal *mphB*, highly homologous to *mphA*, which may explain this phenotype **(Table S4).** Additional multidrug efflux exporters that could modulate susceptibility but do not directly inactivate macrolides were also detected (<u>206</u>). Consistent with this interpretation, the WG5 transconjugant carrying pUTAK-22.1-MDR, which encodes truncated *mphA*, remained susceptible to ERY (data not shown).

### Multiplasmid resistance complement and dosage effects

Plasmid pUTAK-22.2-MDR contributed resistance to five antibiotic classes, including phenicols such as chloramphenicol (CHL; 1× *floR*) (<u>207</u>), tetracycline (TET; 1× *tet(A)*) (<u>208</u>), aminoglycosides such as streptomycin (STR; 1× *aph*(*6*)*-Id*-*aph(3′)-Ib*) (<u>181</u>), sulfonamides such as sulfisoxazole (SFX; 1× *sul2*) (<u>209</u>), complementing the two *sul1* copies on pUTAK-22.1-MDR. The plasmid also carries a β-lactamase gene conferring ampicillin resistance (AMP; 1× *blaCMY-2*) (<u>210</u>) (Fig. 2B). The virulence plasmid pUTAK-22.4-pO111 contributed an additional copy of the *aph*(*6*)*-Id-aph(3′)-Ib* aminoglycoside-resistance module, further increasing streptomycin resistance. Consistent with this ARG inventory across all plasmids, antimicrobial susceptibility testing confirmed resistance to STR, GEN, SFX, KAN, TMP, ERY, CHL, TET, and AMP **(Fig. S3)**. The distribution and redundancy of aminoglycoside resistance genes suggested gene-dosage effects, particularly for STR resistance, supported by multiple copies of the *aph*(*6*)*-Id-aph(3′)-Ib* module and *aadA2.* Similarly, sulfonamide resistance was supported by duplicated *sul* loci. To assess the contribution of each plasmid, STR resistance in UTAK-22 was compared with resistance in WG5 transconjugants carrying various plasmid combinations. Plasmid-linked phenotypes were evaluated using antibiotic-selective readouts in WG5 backgrounds representing subsets of the native plasmid complement. In this system, the wild-type (WT) phenotype reflected the natural plasmid composition of UTAK-22-, whereas individual WG5 transconjugants captured specific plasmid combinations. Streptomycin resistance was provided by a single *aph*(*6*)*-Id*-*aph(3′)-Ib* locus on both pUTAK-22.2-MDR and pUTAK-22.4-pO111 (**Fig. 2B and D**), together with two independent copies of *aadA2* on pUTAK-22.1-MDR **(Fig. 2A)**. Among the backgrounds tested, UTAK-22 exhibited the highest STR tolerance, sustaining growth up to 1024 µg/mL **(Fig. 7)**. The WG5+pUTAK-22.1-MDR/pUTAK-22.2-MDR transconjugant also grew at 1024 µg/mL, but with delayed growth and lower overall OD, whereas WG5+pUTAK-22.1-MDR grew only up to 256 µg/mL (**Fig. S4**). Together, these findings supported a gene-dosage effect in which the combined presence of multiple STR^R^ determinants increased both resistance level and growth tolerance under drug pressure.

**FIG 7.**
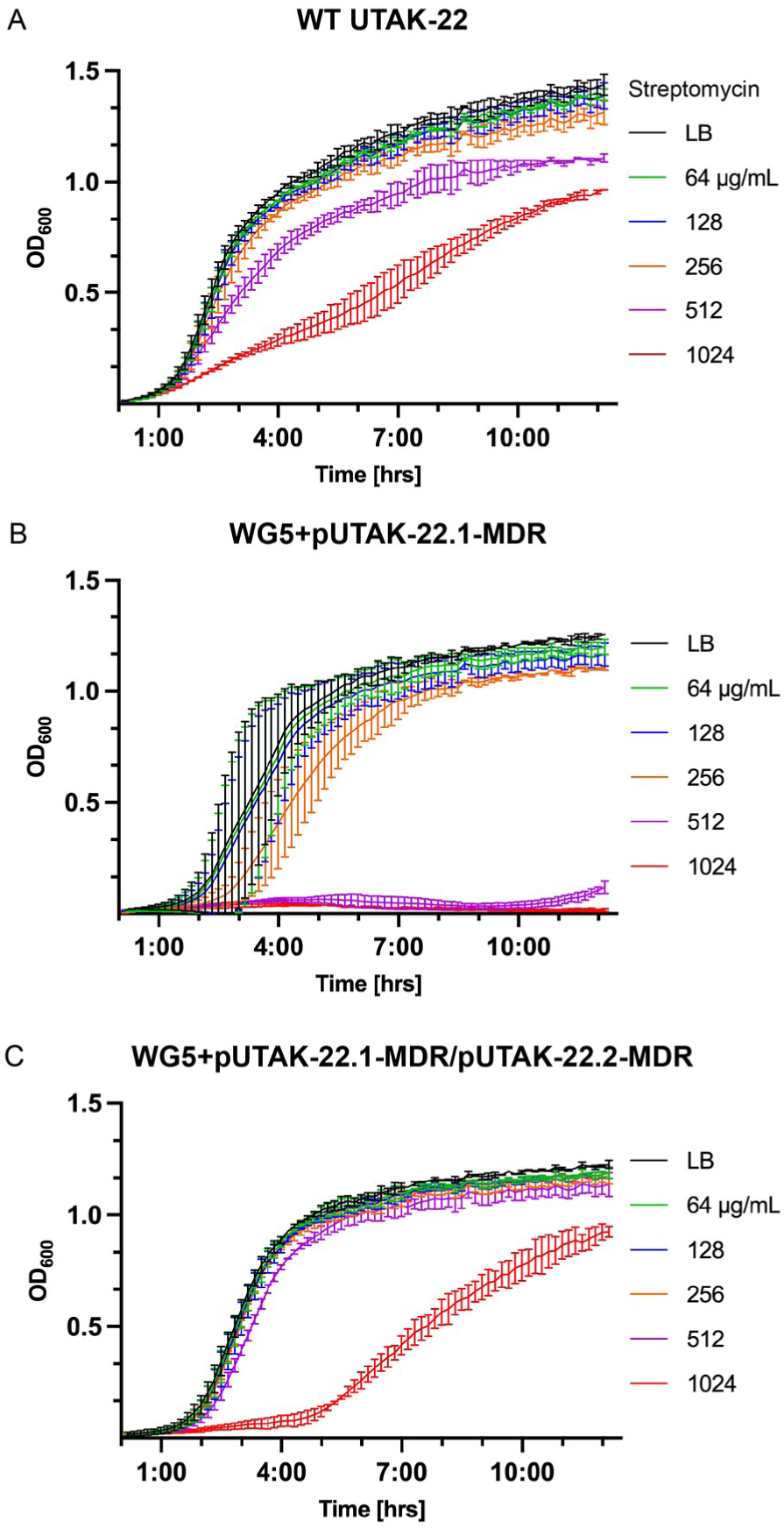
Differential streptomycin tolerance of UTAK-22 and selected WG5 transconjugants. Growth curves of **(A)** UTAK-22, (B) WG5+pUTAK-22.1-MDR, and (C) WG5 carrying the pUTAK-22.1-MDR/pUTAK-22.2-MDR plasmid combination in LB containing increasing streptomycin concentrations (64–1024 µg/mL), monitored by OD₆₀₀ over time relative to drug-free LB controls. WT UTAK-22 sustained growth across all tested concentrations despite dose-dependent reductions in growth kinetics at higher streptomycin levels. WG5+pUTAK-22.1-MDR showed markedly reduced tolerance, whereas the pUTAK-22.1-MDR/pUTAK-22.2-MDR plasmid combination partially restored growth at high streptomycin concentrations. Together, these data support a dosage-dependent contribution of multiple streptomycin-resistance determinants to growth under drug stress. Data are shown as mean ± SD.

## Conclusion

Whole-genome sequencing (WGS) and comparative genomics resolved the chromosome-plasmid architecture of MDR-STEC O111:H8 strain UTAK-22 and revealed a multipartite plasmid system that combined virulence-associated loci with redundant antimicrobial resistance determinants. These data support a model in which pUTAK-22.1-MDR is self-transmissible and can mobilize co-resident plasmids *in trans*, linking resistance dissemination with a broader virulence plasmid network that antibiotic selection can favor (<u>32</u>, <u>65</u>, <u>66</u>, <u>171</u>). The coexistence of overlapping resistance determinants across multiple replicons, together with redundant hemolysin- and streptomycin-associated loci, indicates that plasmid redundancy can strengthen fitness and survival in the native strain background (<u>30</u>, <u>211</u>). Phenotypic comparisons between WT UTAK-22 and selected WG5 transconjugants further supported a dosage-dependent contribution to streptomycin tolerance, while emphasizing that the phenotypes arose from the natural plasmid composition of the strain rather than from the isolated plasmids. The efficient transfer of such plasmids could occur without a major biological cost, which reinforces their persistence (<u>29</u>). Treatment of STEC infections is generally contraindicated because certain antimicrobials, including metronidazole, ampicillin, and trimethoprim-sulfamethoxazole, can induce the SOS response and increase Stx production (<u>96</u>). Macrolides (e.g., azithromycin and erythromycin), rifamycins (e.g., rifampicin), aminoglycosides (e.g., gentamicin and kanamycin), and tetracyclines (e.g., doxycycline) have been shown *in vitro* to limit or avoid Stx release (<u>96</u>). However, MDR among STEC, such as in UTAK-22, narrows an already limited range of usable antibiotics (<u>56</u>): UTAK-22 confers resistance to seven antibiotic classes, β-lactams, trimethoprim, phenicols, sulfonamides, aminoglycosides, tetracyclines, and macrolides. Macrolide resistance is of particular concern, as azithromycin is among the few antibiotics with demonstrated clinical potential (<u>70</u>, <u>212</u>, <u>213</u>). Moreover, resistance to tetracycline and streptomycin is prevalent among food-animal *E. coli*, frequently within MDR profiles (<u>71</u>). Continued genomic surveillance of these hybrid pathogenicity-resistance plasmid systems will be important for tracking dissemination and resistance emergence in O111 STEC lineages (<u>59</u>, <u>64</u>, <u>168</u>).

## Supplemental Figures

**FIG S1.**
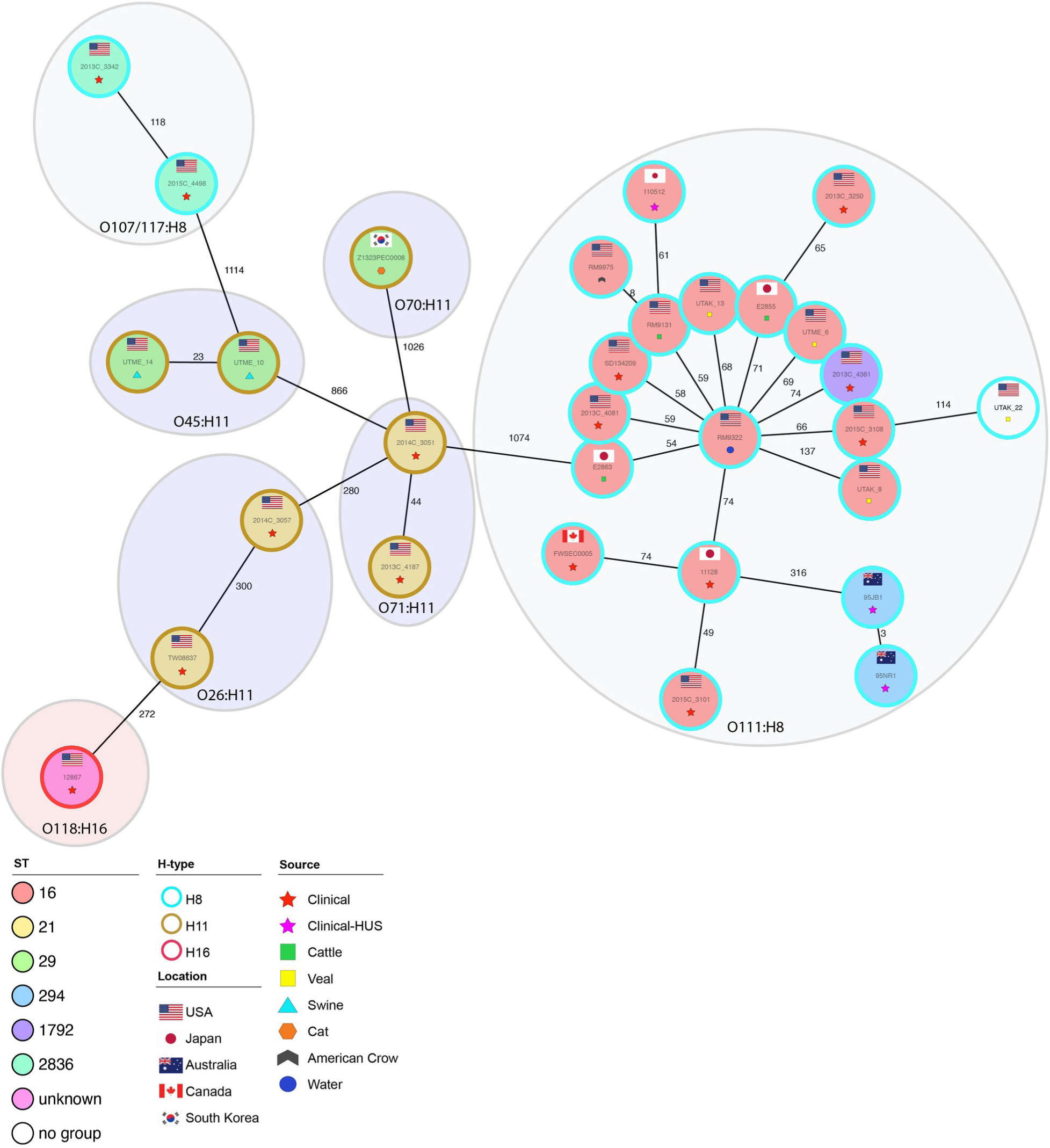
Core-genome phylogeny of UTAK-22 and related *E. coli* genomes. The phylogenetic position of UTAK-22 and the 30 most closely related *E. coli* genomes identified by Mash-based similarity analysis was inferred using a cgMLST minimum-spanning tree generated in Ridom SeqSphere+. The shared gene inventory comprises 4,160 genes, including 3,190 core and 970 accessory loci. Allelic distances between nodes are indicated on the connecting lines. Node color and ring annotations denote sequence type, H-antigen type, source, and geographic origin as indicated in the legend.

**FIG S2.**
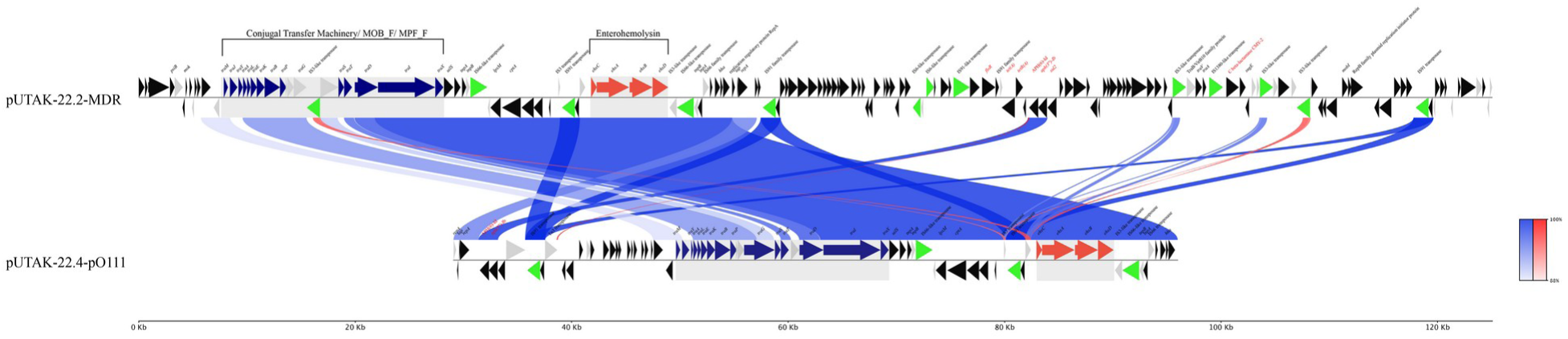
Shared synteny between pUTAK-22.2-MDR and pUTAK-22.4-pO111. Pairwise comparative alignment of pUTAK-22.2-MDR and pUTAK-22.4-pO111 showing extensive conservation across the IncF-like backbone, including replication/stability regions, the shared *ehxCABD* locus, and transfer-associated loci. Shaded links indicate nucleotide similarity between homologous regions. This comparison highlights the close structural relationship between the two plasmids while also showing accessory regions unique to each backbone, including multidrug-resistance cargo in pUTAK-22.2-MDR.

**FIG S3.**
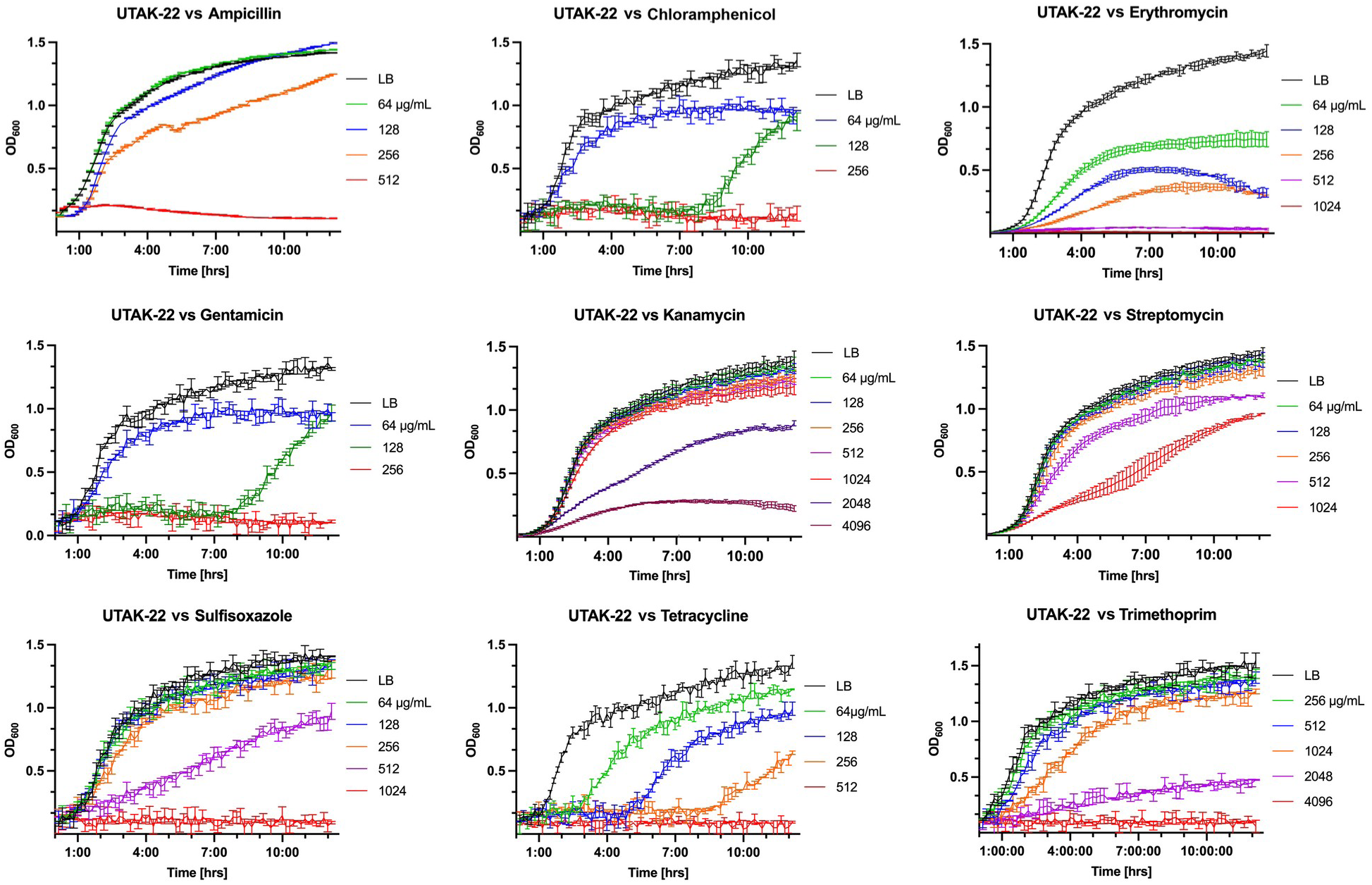
Growth of UTAK-22 across multiple antibiotic conditions. Growth curves of UTAK-22 in LB containing increasing concentrations of ampicillin (AMP), chloramphenicol (CHL), erythromycin (ERY), gentamicin (GEN), kanamycin (KAN), streptomycin (STR), sulfisoxazole (SFX), tetracycline (TET), and trimethoprim (TMP), monitored by OD₆₀₀ over time relative to drug-free LB controls. These plots summarize the antibiotic-response phenotype of the native UTAK-22 plasmid background across multiple drug classes.

**FIG S4.**
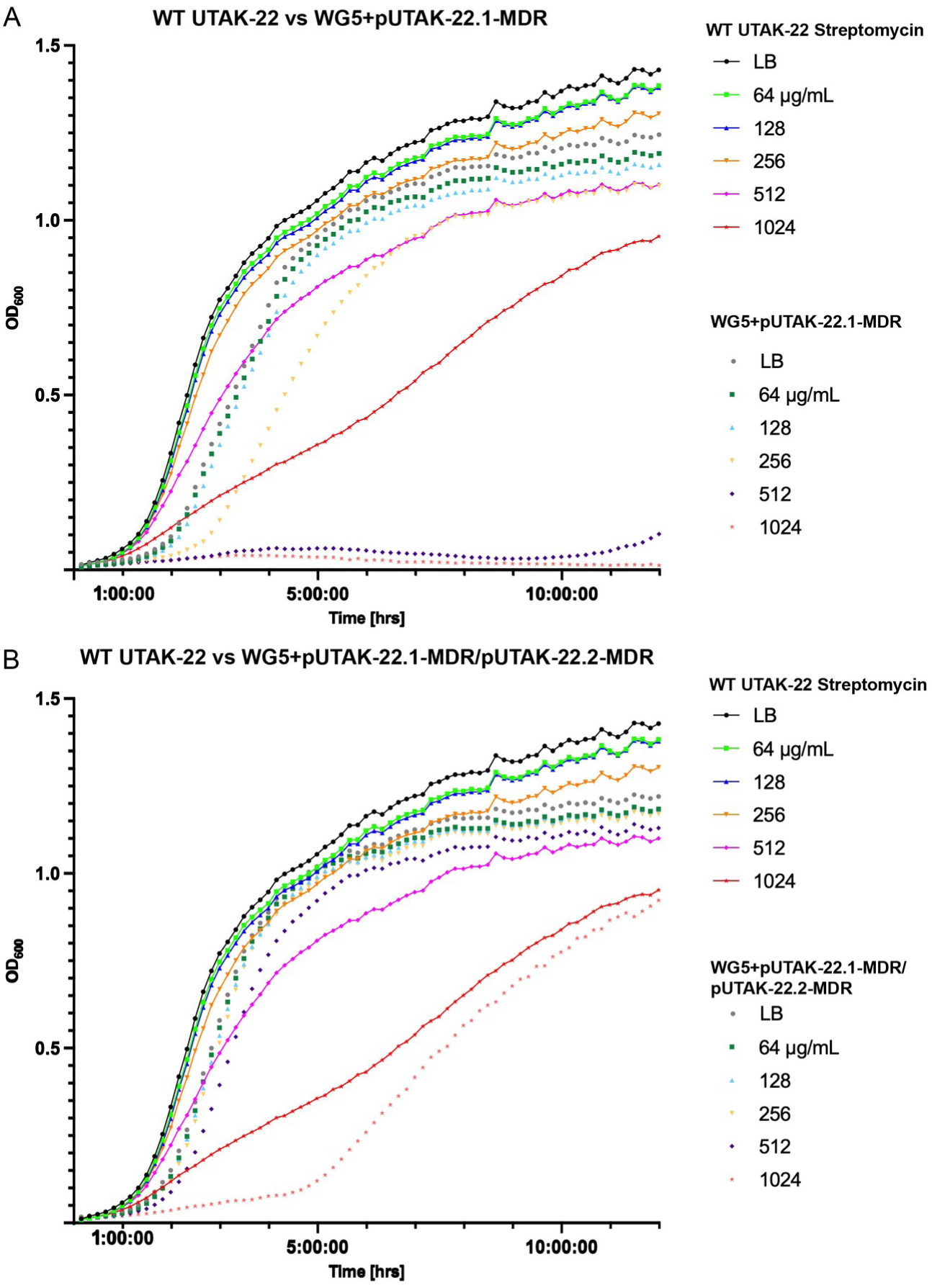
Direct comparison of streptomycin growth kinetics between UTAK-22 and selected transconjugants. Overlay growth curves comparing wild-type UTAK-22 with **(A)** WG5+pUTAK-22.1-MDR and **(B)** WG5 carrying the pUTAK-22.1-MDR/pUTAK-22.2-MDR plasmid combination across increasing streptomycin concentrations. Solid and dotted traces distinguish strain backgrounds as shown in the panel legends. The plots emphasize the higher streptomycin tolerance of the native UTAK-22 plasmid background and the partial restoration of high-dose growth in the co-transfer background relative to pUTAK-22.1-MDR alone.

## Supplemental Tables

Table S1 **Strain-associated metadata and genome statistics**

Table S2 **Primers used in this study**

Table S3 **Achtman 7-gene MLST profiles and cgMLST allele calls (4,160 targets) for UTAK-22 and the 30 most closely related *E. coli* genomes**

Table S4 **Chromosome features**

Table S5 **Plasmid features**

Table S6 **Conjugation efficiency**

## Supplemental Text

Evolutionary reconstruction of MDR cassette dynamics

## Data Availability Statement

The complete genome of *E. coli* serotype O111:H8 strain UTAK-22 has been deposited under BioProject PRJNA1100578. Accessions for underlying Nanopore reads and the assembled and annotated molecules, comprising the chromosome and four carried plasmids, along with strain-associated metadata, are provided in **Table S1**. Supplementary tables, figures, and text are provided as supplementary material accompanying this preprint

## Author contributions

Conceived and designed the experiments: JMB and ME… Analyzed the data: IR, AAK, SSK, MSG, JAA, JMB, and ME. Contributed bioinformatic analysis tools: JAA and ME. ME and SSK developed the PlasmidTyper software used for plasmid taxonomy, mobility, and gene-inventory typing. Performed the conjugation experiments: IR. Assisted with bioinformatics profiling and data visualization: JAA. Provided computational resources: ALR. Drafted the manuscript: ME, IR, and SSK. Contributed to and edited the manuscript: AAK and JMB.

## Acknowledgments

The authors acknowledge the Research Computing Support Group (RCSG) at UT San Antonio for providing computational and high-performance computing resources on the Advanced Research Computing (ARC) cluster that contributed to the reported research. The use of product and company names is necessary to accurately report the methods and results; however, the United States Department of Agriculture (USDA) neither guarantees nor warrants the standard of the products, and the use of names by the USDA implies no approval of the product to the exclusion of others that may also be suitable. The USDA is an equal opportunity provider and employer. We would like to acknowledge Felix Borrego for assistance with data visualization. Mariana Sainz Garcia was supported as an undergraduate McNair Scholar through the Ronald E. McNair Post-Baccalaureate Achievement Program at UTSA.

