## Supplemental Figures for "Pathogenome and Plasmid-Borne Antimicrobial Resistance Phenotypes in a Multidrug-Resistant O111:H8 Shiga Toxin-Producing *Escherichia coli* Strain"

### Figure S1: Core-genome phylogeny of UTAK-22 and related E. coli genomes

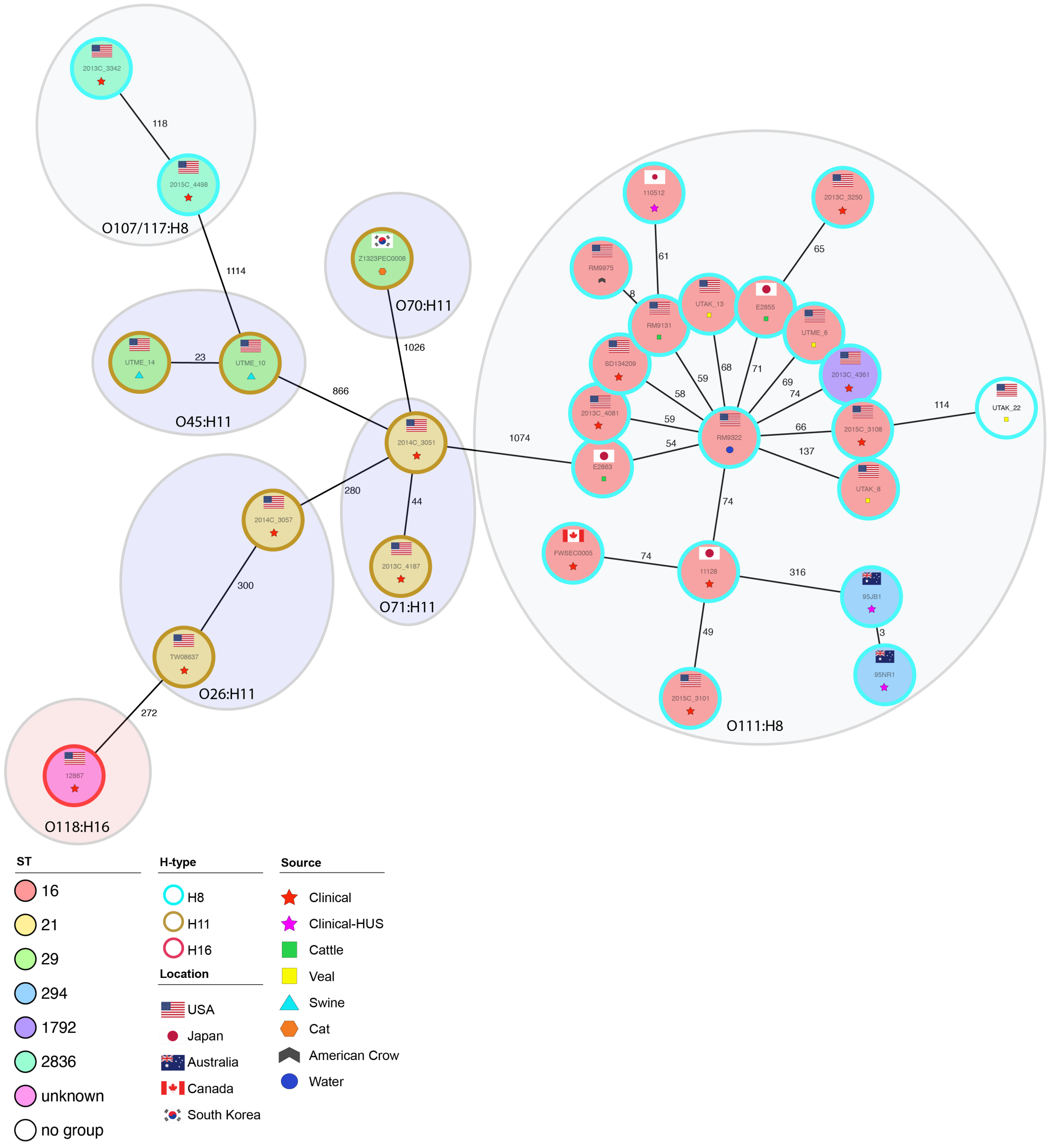


Legend:

Maximum likelihood phylogenetic tree based on concatenated sequence alignments of 2,047 core genes. UTAK-22 is indicated with a filled triangle. Scale bar shows substitutions per site.

### Figure S2: Shared synteny between pUTAK-22.2-MDR and pUTAK-22.4-pO111

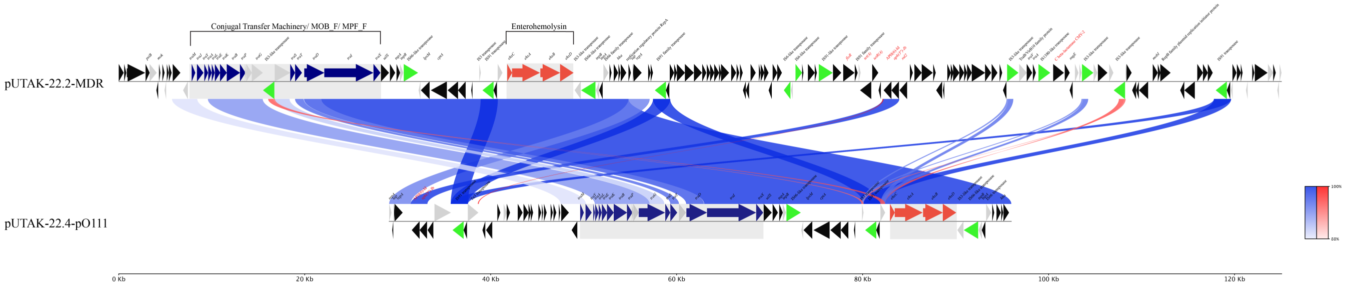


Legend:

BRIG representation of homologous regions between pUTAK-22.2-MDR (innermost ring) and pUTAK-22.4-pO111. Syntenic regions are shown in blue (forward) and red (reverse) orientation.

### Figure S3: Growth of UTAK-22 across multiple antibiotic conditions

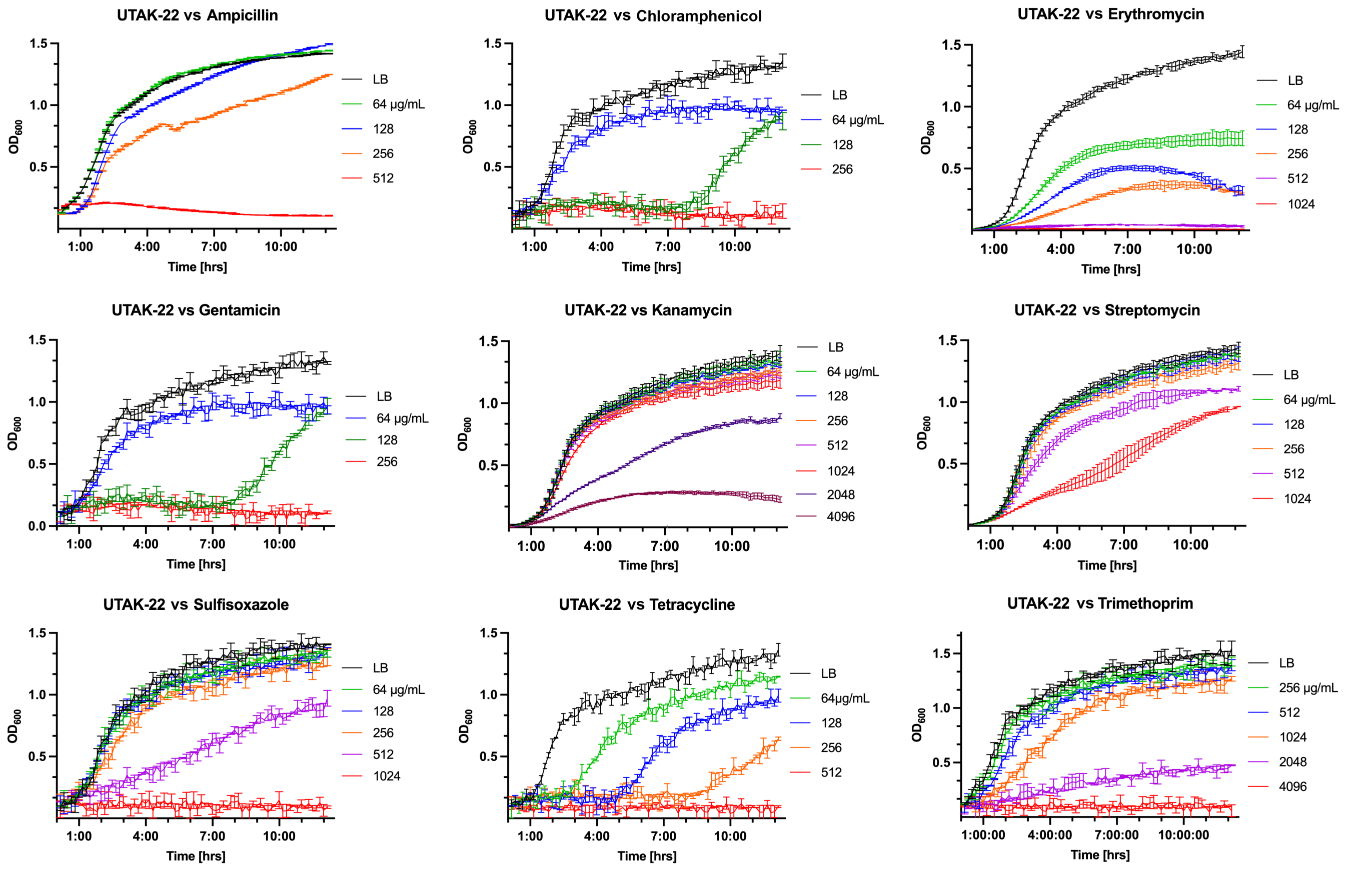


Legend:

Dose-response curves for UTAK-22 grown in liquid culture with increasing concentrations of seven different antibiotics. Each point represents the mean of three biological replicates ± SD.

### Figure S4: Direct comparison of streptomycin growth kinetics between UTAK-22 and se

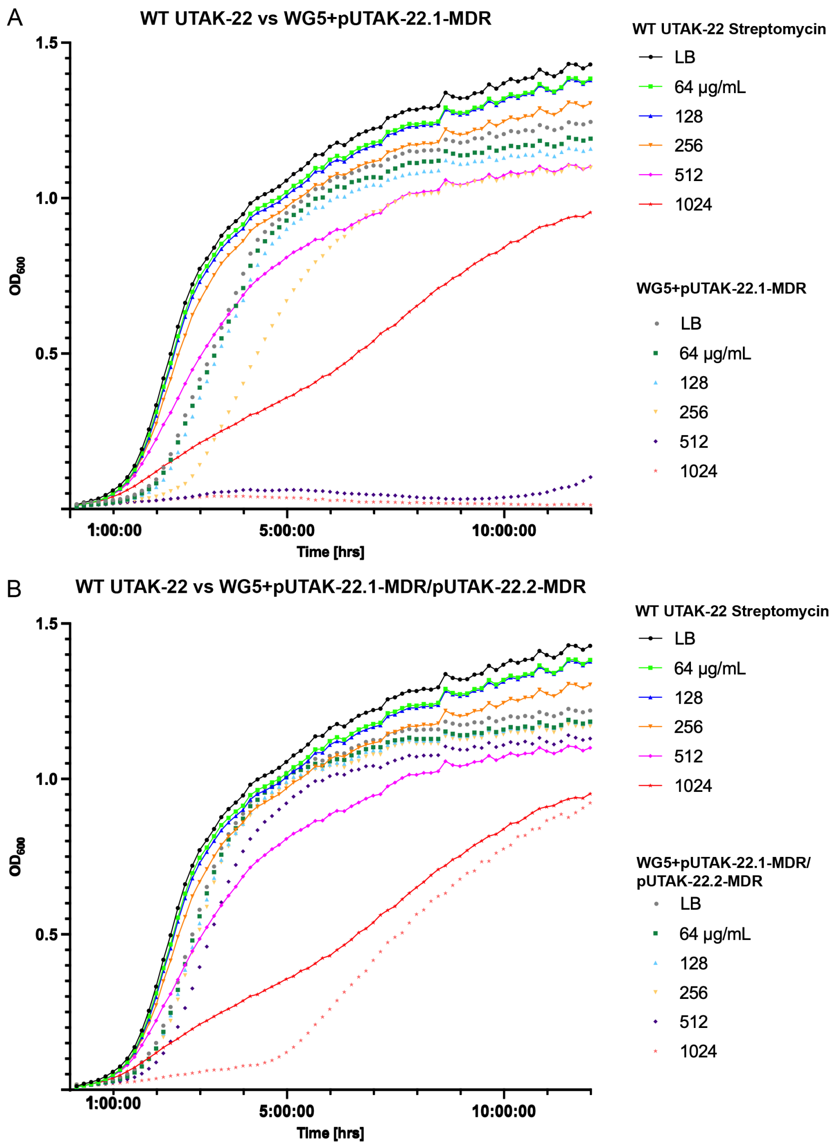


Legend:

Time-series growth curves comparing streptomycin tolerance in UTAK-22 donor and transconjugant strains. OD600 was measured at intervals over 24 hours.
