## Supplemental Text for "Pathogenome and Plasmid-Borne Antimicrobial Resistance Phenotypes in a Multidrug-Resistant O111:H8 Shiga Toxin-Producing *Escherichia coli* Strain"

**Evolutionary Reconstruction of MDR Cassette Dynamics** Class 1 integrons originated within Tn402-like transposons, which provided mobility through the *tni* transposition module (1). Over evolutionary time, many clinically important class 1 integrons have lost all or part of this Tn402 backbone (particularly the *tni* genes), while retaining the core integron platform (*intI1*, *attI1*, *Pc/Pi* promoter, *attC*, and the 3′-conserved segment with *qacEΔ1* and *sul1*). Consistent with this evolutionary trajectory, integron I features both the ancestral Tn402 inner inverted repeat (IRi) and terminal inverted repeat (IRt), whereas integron II retains only IRi; both lack the *tni* transposition genes. The presence of IRi/IRt boundaries supports prior mobilization by a Tn402-like ancestor (2). Consistent with this, both integrons are embedded within a Tn3-like transposon whose *res* site is a known target for Tn402-family insertion (3). Flanking direct repeats were detected only for integron I, marking the insertion site. However, the absence of detectable direct repeats (DR) for integron II is not uncommon, as direct repeats can be obscured by recombination, rearrangement, or decay after integration (4). Class 1 integron I encodes antibiotic resistance genes *aadA* (streptomycin), *aac(3)-VI* (gentamicin), *qacEΔ1* (quaternary ammonium compounds), and *sul1* (sulfonamides), whereas the right class 1 integron encodes *dfrA* (trimethoprim), *aadA* (streptomycin), *qacEΔ1* (quaternary ammonium compounds), *sul1* (sulfonamides), *aph(3′)-Ia* (kanamycin/neomycin), and the inducible macrolide resistance operon *mphR(A)-mrx(A)-mph(A)* (macrolides), with *mph(A)* truncated by IS26 (5).
